# *In vitro* induction and *in vivo* engraftment of lung bud tip progenitor cells derived from human pluripotent stem cells

**DOI:** 10.1101/108845

**Authors:** Alyssa J. Miller, David R. Hill, Melinda S. Nagy, Yoshiro Aoki, Briana R. Dye, Alana M. Chin, Sha Huang, Felix Zhu, Eric S. White, Vibha Lama, Jason R. Spence

## Abstract

The bud tip epithelium of the branching mouse and human lung contains multipotent progenitors that are able to self-renew and give rise to all mature lung epithelial cell types. The current study aimed to understand the developmental signaling cues that regulate bud tip progenitor cells in the human fetal lung, which are present during branching morphogenesis, and to use this information to induce a bud tip progenitor-like population from human pluripotent stem cells (hPSCs) *in vitro*. We identified that FGF7, CHIR-99021 and RA maintained isolated human fetal lung epithelial bud tip progenitor cells in an undifferentiated state *in vitro*, and led to the induction of a 3-dimensional lung-like epithelium from hPSCs. 3-dimensional hPSC-derived lung tissue was initially patterned, with airway-like interior domains and bud tip-like progenitor domains at the periphery. Epithelial bud tip-like domains could be isolated, expanded and maintained as a nearly homogeneous population by serial passaging. Comparisons between human fetal lung epithelial bud tip cells and hPSC-derived bud tip-like cells were carried out using immunostaining, *in situ* hybridization and transcriptome-wide analysis, and revealed that *in vitro* derived tissue was highly similar to native lung. hPSC-derived epithelial bud tip-like structures survived *in vitro* for over 16 weeks, could be easily frozen and thawed and maintained multi-lineage potential. Furthermore, hPSC-derived epithelial bud tip progenitors successfully engrafted in the proximal airways of injured immunocompromised NSG mouse lungs, where they persisted for up to 6 weeks and gave rise to several lung epithelial lineages.

## Introduction

During development, the lung undergoes branching morphogenesis, where a series of stereotyped epithelial bifurcations give rise to the branched, tree-like architecture of the adult lung (Metzger et al., 2008). A population of rapidly proliferating progenitor cells resides at the tips of the epithelium throughout the branching process (‘bud tip progenitors’) (Branchfield et al., 2015; Rawlins et al., 2009). This population, which expresses *Id2* and *Sox9* in mice, has the capability to differentiate into both mature airway and alveolar cell types. At early stages of branching morphogenesis, this population of progenitors gives rise to proximal airway cells, while at later time points these progenitors give rise to alveolar cells (Branchfield et al., 2015; Rawlins et al., 2009).

Studies utilizing genetic mouse models have shown that lung branching morphogenesis and proximal-distal patterning are regulated by a series of complex mesenchymal-epithelial interactions that involve multiple signaling events, transcription factors, and dynamic regulation of the physical environment (Domyan and Sun, 2010; Hines and Sun, 2014; Kim and Nelson, 2012; Morrisey et al., 2013; Morrisey and Hogan, 2010; Rawlins, 2010; Rock and Hogan, 2011; Varner and Nelson, 2014). These studies have identified major roles for several signaling pathways in these processes, including Wnt, Fibroblast Growth Factor (Fgf), Bone Morphogenic Protein (Bmp), Sonic Hedgehog (Shh), Retinoic Acid (RA) and Hippo signaling, among others (Abler et al., 2009; Bellusci et al., 1997a; Bellusci 1997b; Bellusci 1996; Cornett et al., 2013; Desai et al., 2006; Desai 2004; Domyan et al., 2011; Goss et al., 2009; Harris-Johnson et al., 2009; Herriges et al., 2015; Lange et al., 2015; Lu et al., 2009; Mahoney et al., 2014; Motoyama et al., 1998; Sekine et al., 1999; Shu et al., 2005; Weaver et al., 2000; White et al., 2006; Yin et al., 2011; Yin 2008; Zhang et al., 2016; Zhao et al., 2014). However, due to the complex and intertwined nature of these signaling networks, perturbations in one pathway often affect signaling activity of others (Hines and Sun, 2014; Morrisey et al., 2013; Ornitz and Yin, 2012).

These developmental principles, learned from studying model organism development, have been used as a guide to successfully direct differentiation of human pluripotent stem cells into differentiated lung lineages and 3-dimensional lung organoids (Chen et al., 2017; Dye et al., 2016a; Dye 2015; Firth et al., 2014; Ghaedi et al., 2013; Gilpin et al., 2014; Gotoh et al., 2014; Huang et al., 2013; Konishi et al., 2015; Longmire et al., 2012; McCauley et al., 2017). However, specifically inducing and maintaining the epithelial bud tip progenitor cell population from hPSCs has remained elusive. For example, our own studies have shown that hPSCs can be differentiated into human lung organoids (HLOs) that possess airway-like epithelial structures and alveolar cell types; however, it was not clear if HLOs passed through a bud tip progenitor-like stage, mimicking all stages of normal development *in vivo* (Dye et al. 2015). More recent evidence from others has demonstrated that putative bud tip progenitor cells may be induced from hPSCs; however, these cells were rare and were not assessed in detail (Chen et al., 2017). Thus, generation of a robust population of bud tip progenitor cells from hPSCs would shed additional mechanistic light on how these cells are regulated, would provide a platform for further investigation into mechanisms of lung lineage cell fate specification, and would add a layer of control to existing directed differentiation protocols allowing them to pass through this developmentally important progenitor transition.

In the current study, we conducted a low-throughput screen using isolated mouse epithelial bud tip cultures to identify factors that maintained epithelial bud tip progenitors *in vitro*. These conditions were also tested using isolated human fetal epithelial bud tip progenitors cultured *in vitro* and for the ability to induce a bud tip like population from hPSCs. We determined that FGF7 promoted an initial expansion of human epithelial bud tip progenitors, and that the addition of CHIR-99021 (a GSK3β inhibitor that acts to stabilize β-catenin) and All-trans Retinoic Acid (RA) (3-Factor conditions, herein referred to as ‘3F’) were required for growth/expansion of human fetal bud tips as epithelial progenitor organoids that maintained their identity *in vitro*.

When applied to hPSC-derived foregut spheroid cultures, we observed that 3F conditions promoted growth into larger organoid structures with a patterned epithelium that had airway-like and bud tip-like domains. Bud tip-like domains could be preferentially expanded into ‘bud tip organoids’ using serial passaging. hPSC-derived bud tip organoids had a protein expression and transcriptional profile similar to human fetal progenitor organoids. Finally, we demonstrated that hPSC-derived epithelial bud tip organoids can engraft into an injured mouse airway and undergo multilineage differentiation. Taken together, these studies provide an improved mechanistic understanding of human lung bud tip progenitor cell regulation and establish a platform for studying the maintenance and differentiation of lung bud tip progenitor cells.

## Results

### Characterizing human fetal lung bud tip progenitors

In mice, epithelial bud tip progenitor cells express several transcription factors, including *Sox9, Nmyc* and *Id2* (Chang et al., 2013; Moens et al., 1992; Okubo et al., 2005; Perl et al., 2005; Rawlins et al., 2009; Rockich et al., 2013). However, recent studies have suggested that significant differences between murine and human fetal bud tip progenitor cells (Danopoulos et al., 2017; Nikolić et al., 2017). To confirm and extend these recent findings, we carried out an immunohistochemical analysis using well-established protein markers that are present during mouse lung development (Figure 1A-C, Figure S1) on human lungs between 10-20 weeks of gestation. We also conducted RNAsequncing on freshly isolated epithelial lung bud tips, which were dissected to remove mesenchymal cells, to identify genes that were enriched in epithelial progenitors (Figure 1D-E). We note that our approach using manual and enzymatic dissection techniques were unlikely to yield pure epithelial cells, and likely possessed a small population of associated mesenchyme. Consistent with the developing mouse lung (Perl et al., 2005; Rockich et al., 2013), we observed that SOX9 is expressed in bud tip domains of the branching epithelium (Figure 1A, Figure S1A). In contrast to the developing murine lung, we observed SOX2 expression in these bud tip progenitor domains until 16 weeks of gestation, at which time SOX2 expression was lost from this population (Figure 1A, Figure S1A). We also observed expression of *ID2* by *in situ* hybridization (Figure 1B, Figure S1F), with expression becoming increasingly intense in the bud tips as branching progressed, up through 20 weeks gestation (Figure S1F). Bud tips expressed nearly undetectable levels of Pro-SFTPC at 10 and 12 weeks, with low levels of expression detected by 14 weeks (Figure S1D). Pro-SFTPC expression became more robust by 15 and 16 weeks and continued to increase in the bud tips until 20 weeks (Figure 1C; Figure S1D). SOX9+ bud tip cells were negative for several other lung epithelial markers including SFTPB, PDPN, RAGE and HOPX (Figure 1C, Figure S1C-E). We also examined expression of several proximal airway markers, including P63, acetylated-Tubulin (AC-TUB), FOXJ1, SCGB1A1 and MUC5AC and noted that expression was absent from the epithelial bud tip progenitors (negative staining data not shown). We did observe that PDPN and HOPX were expressed in the transition zone/small airways directly adjacent to the SOX9+ bud tip domain at all time points examined (10-20 weeks of gestation) but that this region did not begin to express the AECI maker RAGE until 16 weeks of gestation (Figure 1C; Figure S1 B,C,E). RNA-sequencing (RNAseq) of isolated, uncultured human fetal epithelial lung bud tips (n=2; 59 days, 89 days gestation) supported protein staining analysis of human fetal buds. Differential expression analysis to identify genes enriched in the human fetal bud tips (isolated human fetal epithelial bud tips vs. whole adult lung) identified 7,166 genes that were differentially expressed (adjusted P-value < 0.01; Figure 1D). A curated heatmap highlights genes corresponding to Figure 1A-C and previously established markers of lung epithelial cells (Figure 1E). Human fetal bud tips have recently been shown to have enrichment for 37 transcription factors (Nikolić et al., 2017). In total, 20 of these 37 transcription factors were also enriched in our analysis (Figure 1F), and gene set enrichment analysis (GSEA) confirmed that this enrichment was highly significant (NES = − 1.8, adjusted P-value=9.1e-5). Combined, this data provided a profile of the protein and gene expression in human fetal lung buds prior to 15 weeks gestation (summarized in Figure 1G).

**Figure 1.**
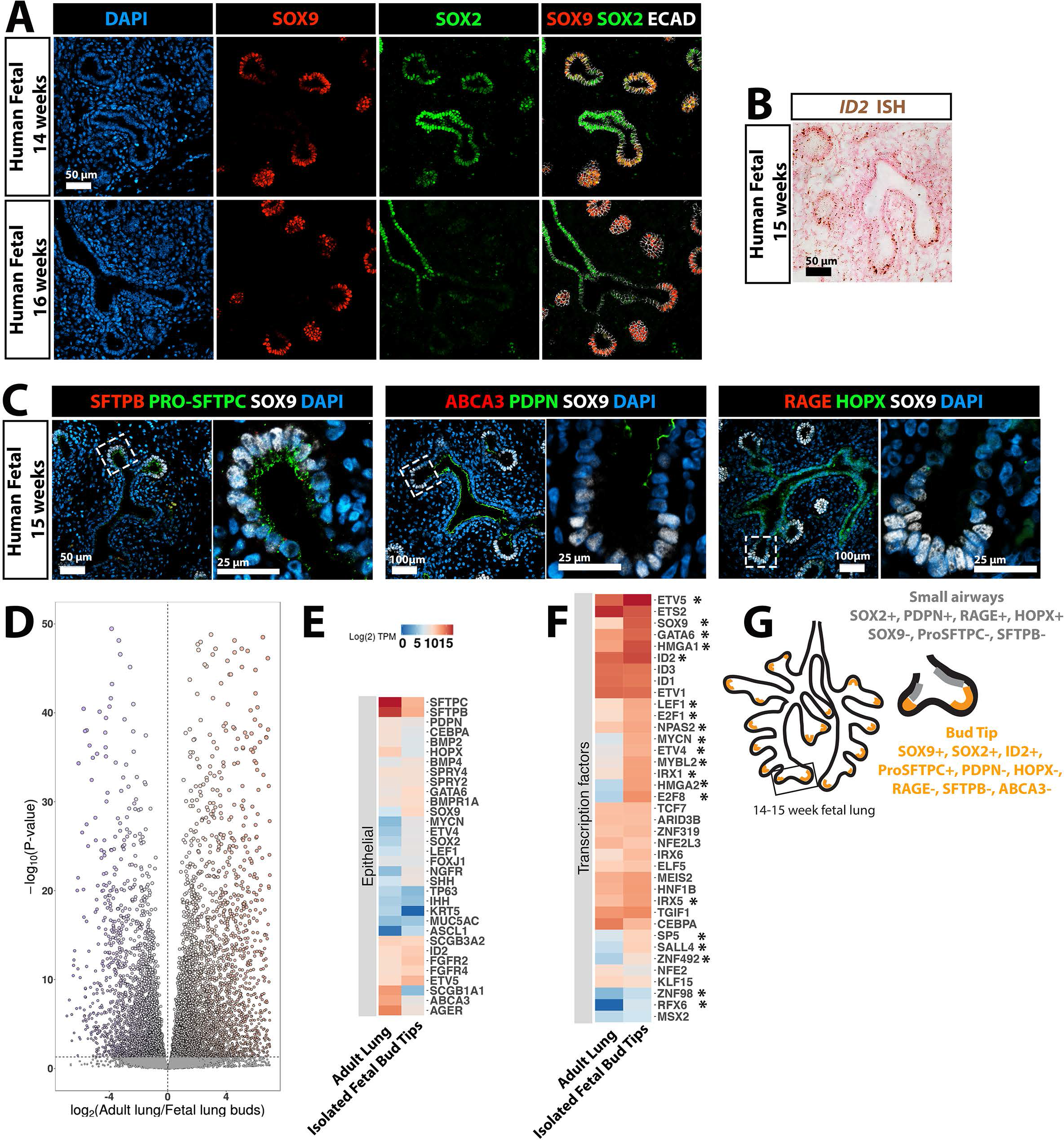
Characterization of bud tip progenitors from 14-16 weeks of human fetal lung development. **(A)** Expression of SOX2 and SOX9 in human fetal lungs at 14 and 16 weeks of gestation. Scale bar represents 50 μm. **(B)** Expression of ID2 in human fetal lungs at 15 weeks gestation as identified by *in situ* hybridization. Scale bar represents 50 μm. **(C)** Expression of SOX9 along with Pro-SFTPC, SFTPB, PDPN,HOPX, ABCA3 or RAGE in human fetal lung at 15 weeks. Scale bars represent 50μm (low magnification images) and 25 μm (high magnification images). **(D)** Volcano plot of differentially expressed identified by comparing isolated, uncultured human fetal bud tips with whole adult human lung. A total of 7,166 genes were differentially expressed (adjusted P-value < 0.01). **(E)** Heatmap showing expression of genes known to be expressed lung epithelial cells in isolated human fetal bud tips and whole adult human lung. **(F)** Heatmap showing expression of 37 human bud-tip enriched transcription factors (Nikolić et al., 2017) in isolated human fetal bud tips and whole adult human lung. 20 of the 37 transcription factors (marked with asterisk “*”) were statistically significantly enriched in isolated fetal bud tips. **(G)** Summary of markers expressed by bud tip cells in regions adjacent to the bud tips at 14-15 weeks gestation as identified by protein staining and *in situ* hybridization.

### Murine bud tip growth *in vitro*

In order to establish an experimental framework that would allow us to efficiently work with rare/precious human tissue samples, we first conducted a low-throughput screen using mouse epithelial bud tips to identify factors that promoted tissue expansion and maintenance of SOX9 expression. Epithelial bud tips were isolated from lungs of embryonic day (E) 13.5 Sox9-eGFP mice and cultured in a Matrigel droplet (Figure S2A). Isolated Sox9-eGFP lung bud tips were shown to express GFP and to have the same level of *Sox9* mRNA as their wild type (WT) counterparts by QRT-PCR analysis (Figure S2B,C). Treating isolated E13.5 mouse bud tips with no growth factors (basal medium control) or individual factors identified from the literature as important for lung development showed that FGF7 robustly promoted growth, expansion and survival of isolated buds for up to two weeks (Figure S1D). Interestingly, the same concentration of FGF7 and FGF10 had different effects on lung bud outgrowth, an observation that could not be overcome even when buds were exposed to a 50-fold excess of FGF10 (Figure S2E).

### FGF7, CHIR-99021 and RA are sufficient to maintain the expression of SOX9 *in vitro*

A lineage trace utilizing isolated epithelial bud tips from Sox9-Cre^ER^;Rosa26^Tomato^ mice showed that FGF7 promoted outgrowth of *Sox9+* distal progenitors cells (Figure S2F). However, *Sox9* mRNA and protein expression were significantly reduced over time (Figure S2G, M).

We therefore sought to identify additional growth factors that could maintain expression of SOX9 *in vitro*. To do this, we grew bud tips in media with all 5 factors included in our initial screen (‘5F’ media), removed one growth factor at a time, and examined the effect on expression of *Sox9* and *Sox2* (Figure S2H-J).

Bud tips were grown in 5F minus one factor for two weeks in culture. Removing any individual factor, with the exception of FGF7, did not affect the ability of isolated buds to expand (Figure S2H). QRT-PCR analysis showed that removing BMP4 led to a statistically significant increase in *Sox9* mRNA expression levels when compared to other culture conditions (Figure S2I). Removing any other single factor did not lead to statically significant changes in *Sox9* expression (Figure S2I). *Sox2* gene expression was generally low in all culture conditions (Figure S2J).

Our data demonstrated that FGF7 is critical for *in vitro* expansion of isolated murine bud tips and removing BMP4 enhanced *Sox9* expression. We therefore screened combinations of the remaining factors to determine a minimal set of factors that could maintain high SOX9 expression (Figure S2K-N). Cultured buds treated with 4-factor ‘4F’(‘4F’ FGF7, FGF10, CHIR-99021, RA) or 3-Factor conditions (‘3F’ FGF7, CHIR-99021, RA) supported the most robust SOX9 protein and mRNA expression (Figure S2L-N)., with no significant differences between SOX9 expression between these two groups. Therefore, a minimum set of 3 factors (FGF7, CHIR-99021, RA) are sufficient to allow growth of mouse epithelial bud tip progenitor cells and to maintain SOX9 expression *in vitro*.

### *In vitro* growth and maintenance of human fetal distal epithelial lung progenitors

We asked if conditions supporting mouse bud tip progenitors also supported growth and expansion of human bud tip progenitors *in vitro*. Distal epithelial lung buds were enzymatically and mechanically isolated from the lungs of 3 different biological specimens at 12 weeks of gestation (84-87 days; n=3) and cultured in a Matrigel droplet (Figure 2A-B). When human bud tips were cultured *in vitro*, we observed that FGF7 promoted an initial expansion of tissue *in vitro* after 2 and 4 weeks, but structures began to collapse by 6 weeks in culture (Figure S3A). All other groups tested permitted expansion and survival of buds as “fetal progenitor organoids” for 6 weeks or longer (Figure 2C; Figure S3A). A description of the nomenclature for different tissues/samples/organoids used in this manuscript can be found in Table 1 of the Methods. Human fetal progenitor organoids exposed to 3F or 4F media supported robust expression of the distal progenitor markers *SOX9, SOX2, ID2 and NMYC* (Figure S3B-C). In contrast, culture in only 2 factors (FGF7+CHIR-99021 or FGF7+RA) did not support robust bud tip progenitor marker expression (Figure S3B-C). QRT-PCR also showed that fetal progenitor organoids cultured in 3F or 4F media expressed low levels of the proximal airway markers *P63, FOXJ1* and *SCGB1A1* when compared to FGF7-only conditions (Figure S3D-E).

**Figure 2:**
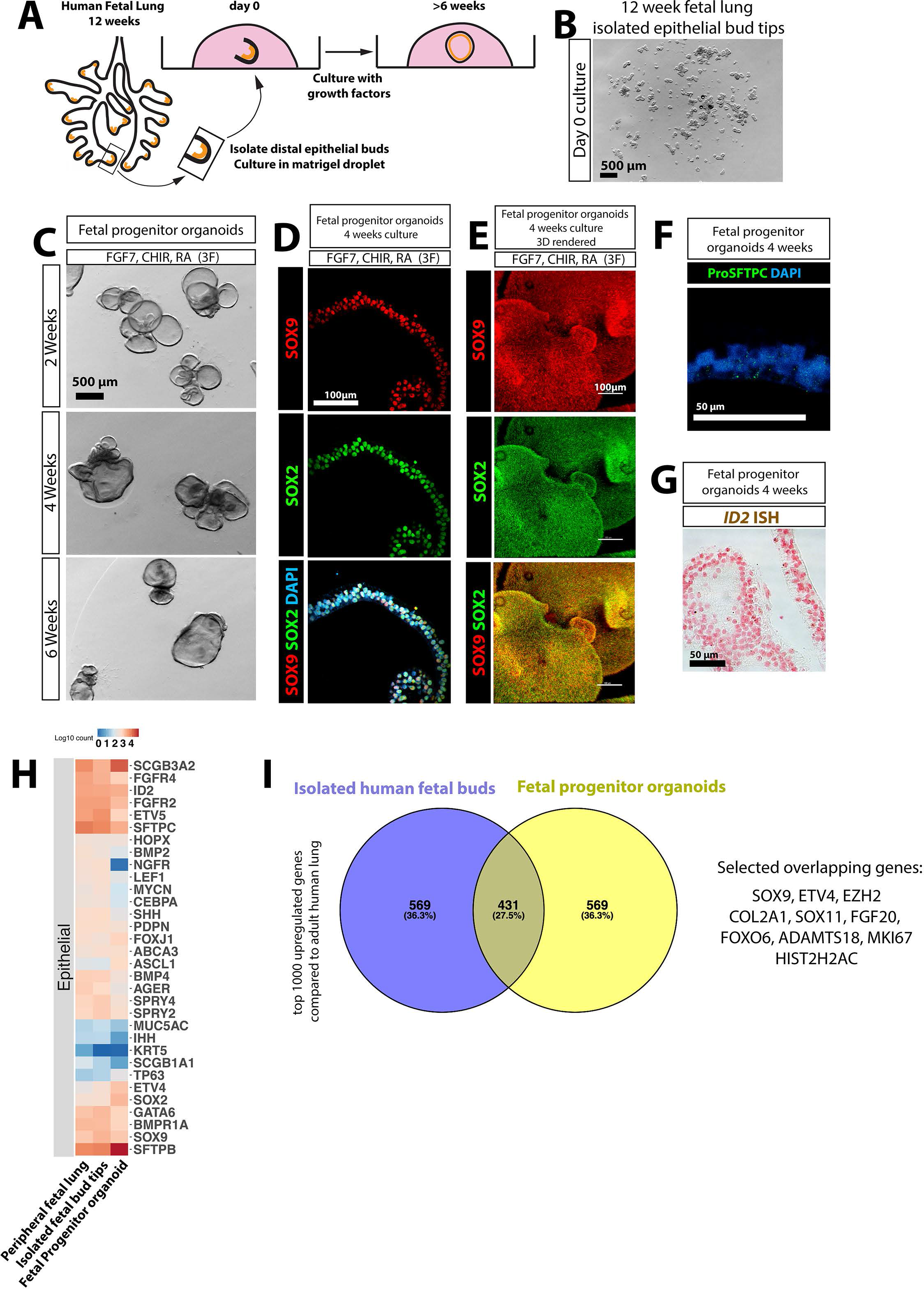
FGF7, CHIR-99021 and RA are sufficient to maintain isolated human fetal bud tip progenitor cells *in vitro*. **(A-B)** Distal epithelial lung bud tips were isolated and cultured in Matrigel droplets. Scale bar in (B) represents 500μm. **(C)** Wholemount brightfield image of human fetal organoids grown in ‘3F’ medium (FGF7, CHIR-99021, RA) at 2, 4 and 6 weeks. Scalebar represents 500 μm. **(D)** Protein staining of SOX2 and SOX9 in sections of fetal progenitor organoids grown in 3F medium. Scale bar represents 100μm. **(E)** Wholemount staining, confocal Z-stack imaging and 3D rendering of SOX2 and SOX9 in fetal progenitor organoids grown in 3F medium. Scale bar represents 100μm. **(F)** Pro-SFTPC after 4 weeks in culture in fetal progenitor organoids. Scale bars represent 50μm. **(G)** *ID2* expression in fetal progenitor organoids after 4 weeks in culture as determined by *in situ* hybridization. Scale bar represents 50μm. **(I)** Heatmap showing expression of genes known to be expressed lung epithelial cells in the whole adult human lung, in isolated human fetal bud tips and in fetal progenitor organoids. **(J)** Differential expression analysis (isolated fetal epithelial bud tips vs. whole adult lung; fetal progenitor organoids vs. whole adult lung) was used to identify the top 1000 most highly upregulated genes from each comparison (log2FoldChange < 0; adjusted p-value < 0.05). Gene overlap was identified using a Venn diagram. 27.5% of genes were common to both groups. A hypergeometric means test showed the number of overlapping genes was highly significant (overlapping p-value=1.4e-278).

**Table 1:** Description of the nomenclature used in this manuscript.

| **Sample nomenclature** | **Description** |
| --- | --- |
| Peripheral fetal lung | The distal/peripheral portion of the fetal lung (i.e. distal 0.5 cm) was excised from the rest of the lung using a scalpel. This includes all components of the lung (e.g. epithelial, mesenchymal, vascular). |
| Isolated fetal bud tip | The bud peripheral portion of the fetal lung was excised with a scalpel and subjected to enzymatic digestion and microdissection. The epithelium was dissected and separated from the mesenchyme, but a small amount of associated mesenchyme likely remained. |
| Fetal progenitor organoid | 3D organoid structures that arose from culturing isolated fetal epithelial bud tips. |
| Foregut spheroid | 3D foregut endoderm structure as described in Dye et al., 2015. Gives rise to PLO when grown in 3F media. |
| Patterned lung organoid (PLO) | Lung organoids that were generated by differentiating hPSCs, as described throughout the manuscript. |
| Bud tip organoid | Organoids derived from PLOs, enriched for SOX9/SOX9 co-expressing cells, and grown/passaged in 3F medium. |

Collectively, our experiments suggested that FGF7, CHIR99021 and RA represent a minimal set of additives required to maintain SOX9+/SOX2+ fetal progenitor organoids *in vitro*. Supporting mRNA expression data, we observed robust SOX9 and SOX2 protein expression as demonstrated by immunofluorescence in sections or by whole mount staining after 4 weeks in culture (Figure 2D-E). Consistent with expression data in lung sections prior to 16 weeks gestation, we observed very weak pro-SFTPC protein expression in human bud tip progenitor organoids. Low levels of *ID2* mRNA were also detected using *in situ* hybridization (Figure 2F-G). Protein staining for several markers was not detected in fetal progenitor organoids treated for 4 weeks in 3F medium, including P63, FOXJ1, SCGB1A1, MUC5AC, HOPX, RAGE, and SFTPB (n=8; negative data not shown), consistent with human fetal epithelial bud tips prior to 16 weeks gestation (Figure S1A-E).

We also performed RNA-sequencing on tissue from the distal portion of fetal lungs (epithelium plus mesenchyme) (n=3, 8, 12 and 18 weeks), on freshly isolated human epithelial bud tips (n=3, 8, 12 and 12 weeks) and on human fetal bud tip progenitor organoids grown in 3F media for 4 weeks in culture (n=2 12 week biological samples, run with statistical triplicates). Analysis revealed a high degree of similarity across samples when comparing epithelial gene expression (Figure 2H). Additionally, we identified genes highly enriched in isolated fetal epithelial bud tips by conducting differential expression analysis on RNAseq data. For this analysis, we compared whole human adult lung versus uncultured isolated lung buds (12 weeks gestation) and versus cultured fetal progenitor organoids (12 weeks gestation, cultured for 2 weeks). In these analyses, whole human lung was used as a ‘baseline’, with the assumption that average expression of any given gene would be low due to the heterogeneous mixture of cells pooled together in the sample, providing a good comparator to identify genes that were enriched in fetal tissue and organoid samples. The top 1000 upregulated genes in bud tips and in fetal progenitor organoids were identified (log2FoldChange < 0; adjusted p-value < 0.05). When comparing upregulated genes, we observed that 431 (27.5%) of the genes were commonly upregulated in both fetal bud tips and cultured fetal organoids, which was highly statistically significant (p-value = 1.4e-278 determined by a hypergeometric test)(Figure 2I).

### FGF7, CHIR-99021, RA induce a bud tip progenitor-like population of cells from hPSC-derived foregut spheroids

Given the robustness by which 3F medium (FGF7, CHIR-99021, RA) supported mouse and human epithelial lung bud tip growth and maintenance *in vitro*, we sought to determine whether these culture conditions could promote an epithelial lung tip progenitor-like population from hPSCs. NKX2.1+ ventral foregut spheroids were generated as previously described (Dye et al., 2016a; Dye 2015), and were cultured in a droplet of Matrigel and overlaid with 3F medium. Spheroids were considered to be “day 0” on the day they were placed in Matrigel (Figure 3A-B). Foregut spheroids cultured in 3F medium generated patterned organoids (hereafter referred to as ‘hPSC-derived patterned lung organoids’ PLOs) that grew robustly, demonstrating predictable growth patterns and gene expression reproducible across several different embryonic and induced pluripotent stem cell lines (n=4) (Figure 3B; Figure S4). PLOs survived for over 16 weeks in culture, and could be frozen, thawed and re-cultured (Figure S4).

**Figure 3:**
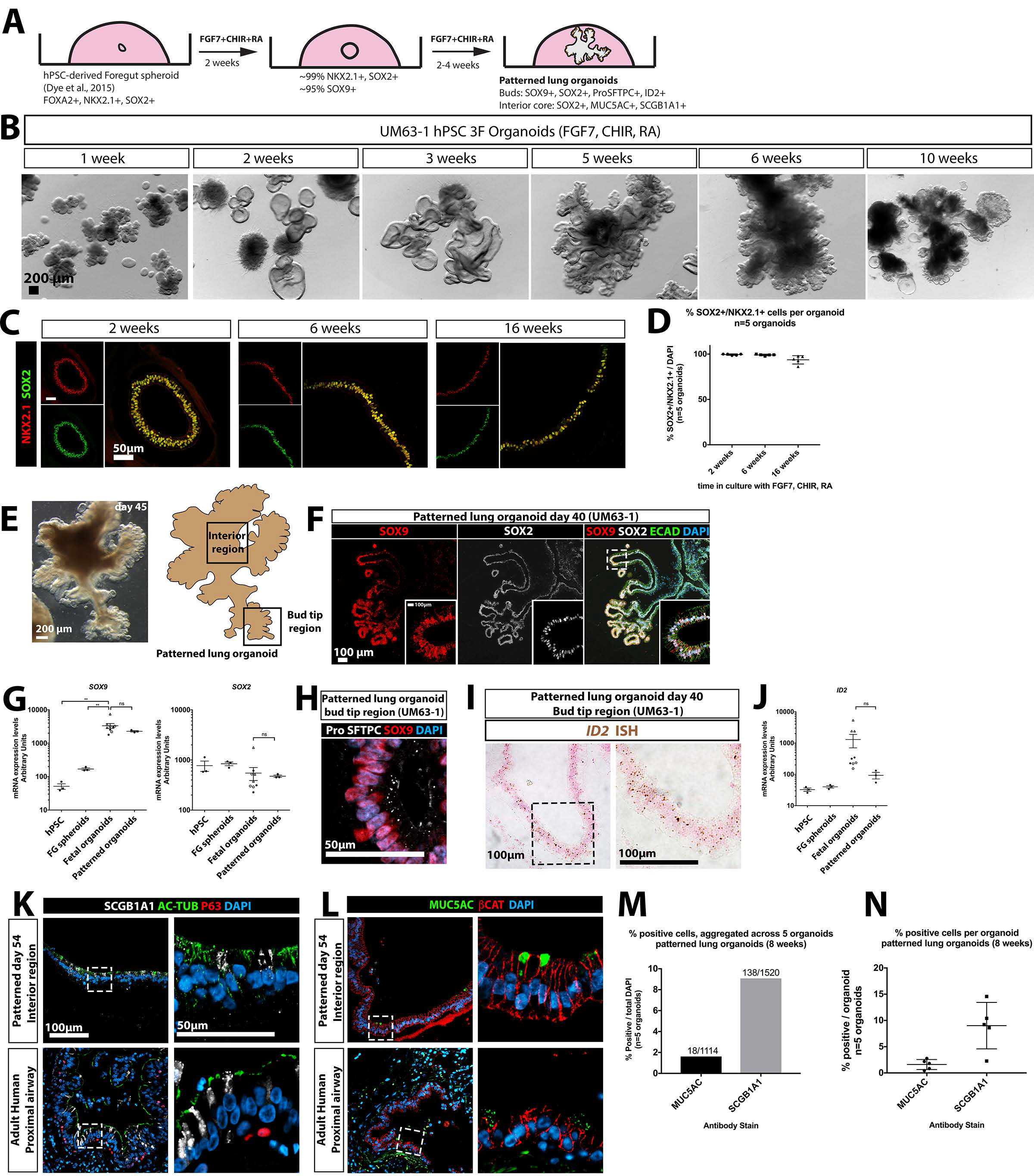
FGF7, CHIR-99021 and RA generate patterned epithelial lung organoids from hPSCs. **(A)** Schematic of approach to derive patterned lung organoids (PLOs) from human pluripotent stem cells. **(B)** Brightfield images of hPSC-derived foregut spheroids cultured in 3F medium (fGf7, CHIR-99021 and RA) and grown *in vitro*. Images taken at 2, 3, 5, 6 and 10 weeks. Scale bars represent 200μm. **(C)** Immunostaining for NKX2.1 and SOX2 in PLOs at 2, 6 and 16 weeks. Scale bars represent 50μm. **(D)** Quantitative analysis of cells co-expressing NKX2.1 and SOX2 in PLOs at 2, 6 and 16 weeks, as shown in (B). Each data point represents an independent biological replicate, and the mean +/− the standard error of the mean is shown for each group. **(E)** Brightfield image of a patterned lung organoid after 6 weeks (45 days) in culture, showing distal budded regions and interior regions (Scale bar represents 200μm) and a schematic representing a patterned lung organoid, highlighting bud tip region and interior regions. **(F)** PLOs co-stained for SOX9 and SOX2 protein expression by immunofluorescence. Inset shows high magnification of boxed region. Scale bar represents 100 μm. **(G)** QRT-PCR analysis of *SOX9* and *SOX2* in undifferentiated hPSCs (H9 hESC line), in foregut spheroids (FG), fetal progenitor organoids and patterned lung organoids. **(H)** Pro-SFTPC and SOX9 co-expression in bud tip region of a PLO. Scale bars represent 50 μm. **(I)** *ID2* expression in PLOs after 40 days *in vitro* as determined by *in situ* hybridization. Scale bar represents 100μm. **(J)** QRT-PCR analysis of *ID2* in undifferentiated hPSCs (H9 hESC line), in foregut spheroids (FG), fetal progenitor organoids and patterned lung organoids. **(K)** Interior regions of patterned lung organoids (top) and adult human airway (bottom) co-stained for SCGB1A1, Acetylated Tubulin (AC-TUB) and P63. Scale bars represent 100μm (left panels, low magnification) or 50μm (right panels, high magnification). **(L)** Interior regions of patterned lung organoids (top) and adult human airway (bottom) co-stained for MUC5AC and the epithelial marker β-catenin (βCAT). Scale bars represent 100 μm (left panels, low magnification) or 50μm (right panels, high magnification). **(M-N)** Percent of cells expressing MUC5AC or SCGB1A1, plotted as aggregate data ((I); # cells positive in all organoids/total cells counted in all organoids) and for each individual patterned lung organoid counted ((J); # cells positive in individual organoid/all cells counted in individual organoid).

At 2 and 6 weeks, PLOs co-expressed NKX2.1 and SOX2 in >99% of all cells (99.41 +/− 0.82% and 99.06 +/− 0.83%, respectively), while 16 week old PLOs possessed 93.7 +/− 4.6% NKX2.1+/SOX2+ cells (Figure 3C-D). However, PLOs at this later time point appeared less healthy (Figure 3B). Interestingly, no mesenchymal cell types were detected in patterned lung organoid cultures at 2, 6 or 16 weeks by immunofluorescence (VIM, α-SMA, PDGFRα; negative immunostaining data not shown), and QRT-PCR analysis confirms very low expression of mesenchymal markers in PLOs generated from n=3 hPSC lines (Figure S4H).

PLOs also exhibited regionalized proximal-like and bud tip-like domains (Figure 3E-F). In 100% of analyzed PLOs (n=8) Peripheral budded regions contained cells that co-stained for SOX9 and SOX2, whereas interior regions of the PLOs contained cells that were positive only for SOX2 (Figure 3E-F). PLOs expressed similar levels of *SOX2* and *SOX9* when compared to fetal progenitor lung organoids (Figure 3G). Budded regions of PLOs also contained SOX9+ cells that weakly co-expressed pro-SFTPC, and *ID2* based on *in situ* hybridization and QRT-PCR (Figure 3H-J).

Interior regions of PLOs contained a small number of cells that showed positive immunostaining for the club cell marker SCGB1A1 (9%) and the goblet cell marker MUC5AC (1%), with similar morphology to adult human proximal airway secretory cells (Figure 3K-N). No multiciliated cells were present, as Acetylated-Tubulin was not localized to cilia and FOXJ1 staining was absent (Figure 2K). Additionally, the basal cell marker P63 was absent from PLOs (Figure 2K), as was staining for markers of lung epithelial cell types including HOPX, RAGE, PDPN, ABCA3, SFTPB, and CHGA (negative data not shown).

### Expansion of epithelial tip progenitor-like cells from patterned lung organoids

PLOs gave rise to epithelial cysts after being passaged by mechanical shearing through a 27-gauge needle, followed by embedding in fresh Matrigel and growth in 3F medium (Figure 4A-B). PLOs were successfully needle passaged as early as 2 weeks and as late as 10 weeks with similar results. Needle passaged cysts can be generated from hPSCs in as little as 24 days (9 days to generate foregut spheroids, 14 days to expand patterned lung organoids, 1 day to establish cysts from needle passaged organoids)(Figure 4A). Needle passaged epithelium reestablished small cysts within 24 hours and could be serially passaged every 7-10 days (Figure 4C). The cysts that formed after needle passaging were NKX2.1+ (Figure S5A) and cells co-expressed SOX2 and SOX9 (Figure 4D-E). Based on these protein staining patterns, we refer to these cysts as ‘bud tip organoids’. When compared to PLOs, bud tip organoids possessed a much higher proportion of SOX9+ cells (42.5% +/− 6.5%, n=5; vs. 88.3% +/− 2.3%, n=9; Figure 4E) and proliferating cells assed by KI67 immunostaining (38.24% +/− 4.7%, n=9 for bud tip organoids vs. 4.9% +/− 0.6%, n=5 for PLOs; Figure 4F-I). In PLOs, we noted that proliferation was largely restricted to SOX9+ budded regions, but only a small proportion of SOX9+ cells were proliferating (8.1% +/− 0. 9%, n=5), whereas bud tip organoids had a much higher proportion of proliferative SOX9+ cells (40.2% +/− 4.3%, n=9) (Figure 4I). Together, this data suggests that needle passaging enriches the highly proliferative bud tip regions of PLOs.

**Figure 4:**
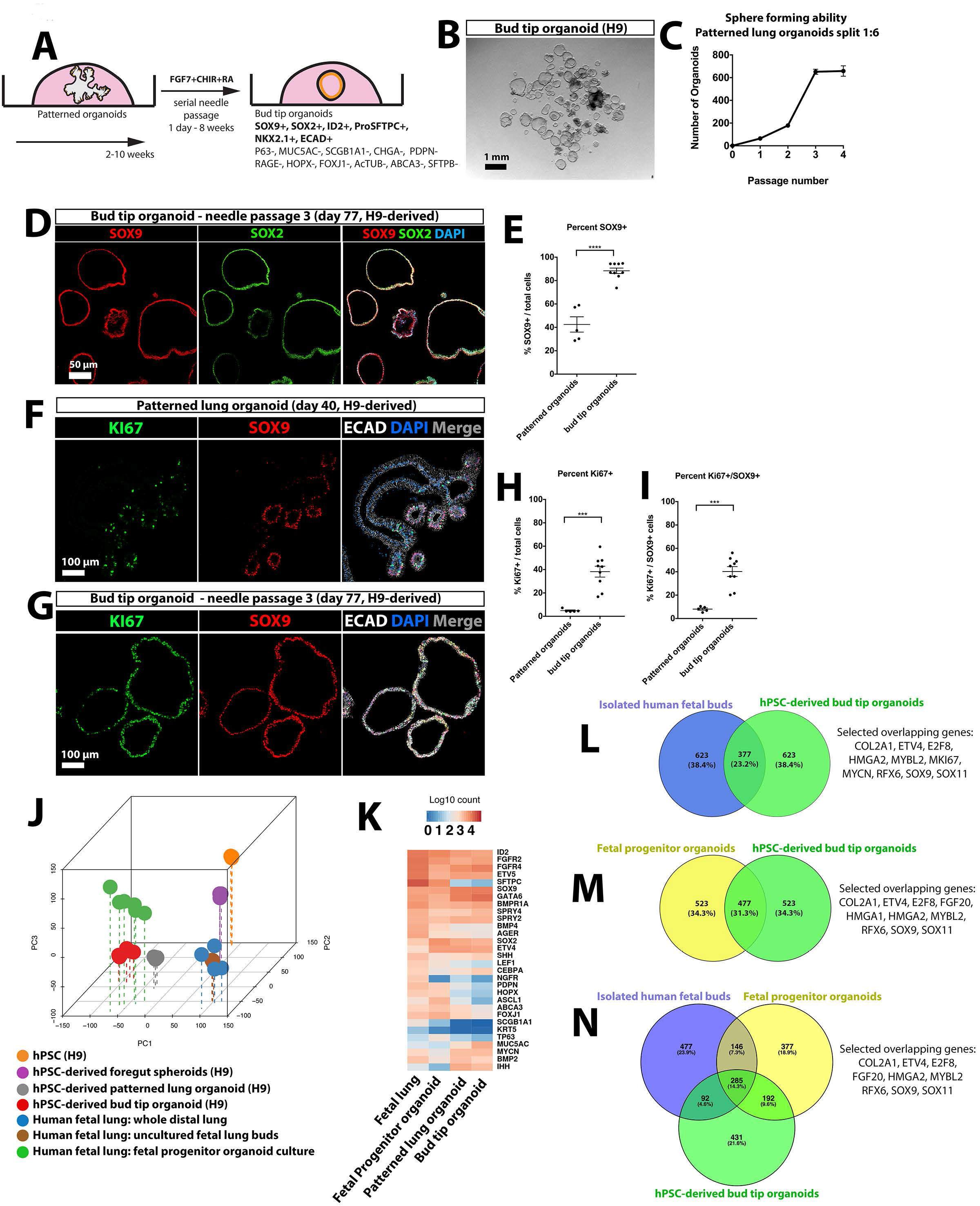
Proliferative SOX9+/SOX2+ progenitors can be expanded from patterned lung organoids. **(A)** Schematic of approach passage PLOs and expand bud tip organoids. **(B)** Needle sheared epithelial fragments were replated in a fresh Matrigel droplet that subsequently formed cystic structures, called ‘bud tip organoids’. Scale bar represents 1mm. **(C)** Quantitative assessment of organoid passaging and expansion. One single patterned lung organoid was needle passaged into 6 wells (passage 1), generating 75 new bud tip organoids in total (average 12.5 per well). A single well of the resulting bud tip organoids were then passaged into 6 new wells after 2 weeks in culture (1:6 split ratio), generating 200 new organoids in total (average 33 per well). This 1:6 passaging was carried out two additional times, every 1-2 weeks for up to 4 passages before growth plateaued. 3 biological replicates were performed for expansion experiments; graph plots the mean +/− the SEM. **(D)** Immunostaining for SOX9 and SOX2 in bud tip organoids. Scale bar represents 50 μm. **(E)** Quantitation of the percent of SOX9+ cells in PLOs and bud tip organoids (# SOX9+ cells/total cells). Each data point represents an independent biological replicate and graphs indicate the mean +/− the standard error of the mean for each experimental group. A two-way unpaired Student’s T test was performed to compare the means of each group. A significance level of 0.05 was used. Significance is shown on the graph according to the following: P > 0.05 ns, P ≤ 0.05 *, P ≤ 0.01 **, P ≤ 0.001 ***, P ≤ 0.0001 ****. **(F-G)** Immunostaining for KI67 and SOX9 in patterned lung organoids (F) and bud tip organoids (G). Scale bar represents 100um. **(H-I) (H)** Quantitation of the percent of all cells that were KI67+ in patterned and bud tip organoids (# KI67+ cells /total cells). **(I)** Quantitation of the percent of proliferating SOX9+ cells in patterned and bud tip organoids (# KI67+/SOX9+ cells/total cells). Each data point represents an independent biological replicate and graphs indicate the mean +/− the standard error of the mean for each experimental group. Significance was determined by an unpaired Student’s T test. A significance value of 0.05 was used. P > 0.05 ns, P ≤ 0.05 *, P ≤ 0.01 **, P ≤ 0.001 ***, P ≤ 0.0001 ****. **(J)** Principal component analysis of RNA-sequencing data to compare the global transcriptome of hPSCs, foregut spheroids, hPSC-derived patterned lung organoids, hPSC-derived bud tip organoids, whole peripheral (distal) fetal lung, freshly isolated (uncultured) fetal lung buds and fetal progenitor organoids. **(K)** Heatmap showing expression of genes known to be expressed in lung epithelial cells in freshly isolated (uncultured) fetal lung buds and fetal progenitor organoids cultured for 2 weeks. **(L-N)** Differential expression analysis of: 1. isolated fetal bud tips vs. whole adult lung; 2. fetal progenitor organoids vs whole adult lung; 3. bud tip organoids vs. whole adult lung was used to identify the top 1000 most highly upregulated genes from each comparison (log2FoldChange < 0; adjusted p-value < 0.05). A Venn diagram illustrates common upregulated genes in **(L)** isolated bud tips and bud tip organoids; **(M)** fetal progenitor organoids and bud tip organoids **(N)** isolated bud tips, fetal progenitor organoids, bud tip organoids. **(L)** hypergeometric means test found that the shared gene overlap was highly significant (overlapping p-value=9.3e-901); **(M)** hypergeometric means test found that the shared gene overlap was highly significant (overlapping p-value=1.2e-1021). **(N)** 285 overlapping genes were shared between the three groups. These genes represented 14.3% of all genes included in the comparison. A small subset of genes previously associated with human or mouse bud-tip progenitor cells are highlighted as “Selected overlapping genes” (L-N).
.

Bud tip organoids were further characterized using *in situ* hybridization and immunofluorescence (Figure S5). Bud tip organoids exhibited protein staining patterns consistent with 12-14 week fetal lungs, including the absence of HOPX, RAGE, PDPN, and ABCA3, while we did observe low expression of *ID2* (Figure S5B-D). Furthermore, no positive protein staining was detected for the proximal airway markers P63, FOXJ1, AC-TUB, MUC5AC, SCGB1A1 or CHGA (negative data not shown).

We then conducted unbiased analysis of bud tip organoids using RNA-sequencing to compare: i) hPSC-derived bud tip organoids; ii) whole peripheral (distal) human fetal lung tissue; iii) freshly isolated fetal bud tips; human fetal lung progenitor organoids; iv) undifferentiated hPSCs; v) hPSC-derived lung spheroids. Principal component analysis (PCA) and Spearman’s correlation clustering revealed the highest degree of similarity between hPSC-derived bud tip organoids, patterned lung organoids and human fetal organoids (Figure 4J; Figure S5F). Interestingly, freshly isolated bud tips and whole distal human fetal lungs clustered together, while all cultured tissues (bud tip organoids, PLOs, fetal progenitor organoids) clustered together, likely reflecting gene expression similarities driven by the tissue culture environment. This analysis highlights a high degree of molecular similarity between human fetal and hPSC-derived organoids grown *in vitro* (Figure 4J).

Differential expression of RNAseq data was also used to interrogate the relationship between tissues. We obtained upregulated genes from the following comparisons: i) uncultured, freshly isolated lung buds (8 and 12 weeks gestation) versus whole adult lung tissue; ii) cultured fetal progenitor organoids (12 weeks gestation, cultured for 2 weeks) versus whole adult lung tissue; iii) hPSC-derived bud tip organoids versus whole adult lung tissue. The top 1000 upregulated genes in each of the three groups were identified (log2FoldChange < 0; adjusted p-value < 0.05), and overlapping genes were identified (Figure 4L-M). A hypergeometric means test found that overlap of enriched genes from isolated human bud tips and hPSC-derived bud tip organoids were highly significant, with 377 (23.2%) overlapping genes (p-value=9.3e-901; Figure 4L). Overlap between cultured fetal progenitor organoids and hPSC-derived bud tip organoids was also highly significant, with 477 (31.3%) overlapping genes (p-value=1.2e-1021) (Figure 4M). Of note, when all three groups were compared, a core group of 285 common upregulated genes were identified representing 14.3% of genes. Of these 285, several have been associated with human or mouse fetal bud tips (Nikolić et al., 2017; Rockich et al., 2013) (Figure 4N; *COL2A1, ETV4, E2F8, FGF20, HMGA2, MYBL2, RFX6, SALL4, SOX9, SOX11*).

### hPSC-derived bud tip organoids maintain multilineage potential *in vitro and in vivo*

In order to demonstrate that hPSC-derived organoids have *bona fide* progenitor potential, we took approaches to differentiate them *in vitro* (Figure 5) and by transplanting them into injured mouse lungs *in vivo* (Figure 6 and Figure S6). For these experiments, we used induced pluripotent stem cells (iPSC) derived foregut spheroids to generate bud tip organoids (iPSC line 20-1 (iPSC20-1); (Spence et al., 2011)).

**Figure 5:**
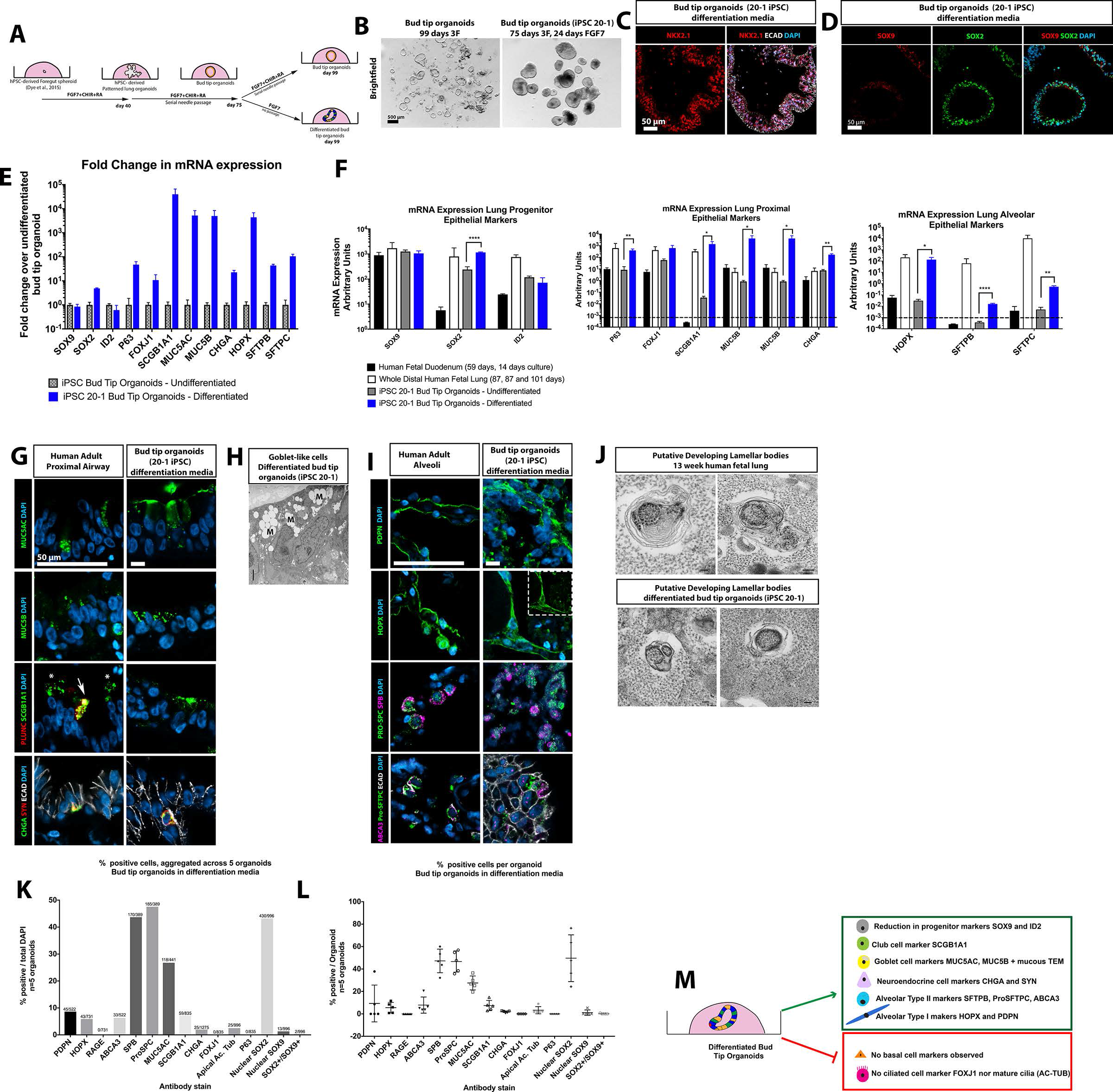
Bud tip organoids retain multilineage potential *in vitro*. **(A)**Schematic of experimental setup. iPSC20-1 bud tip organoids were initially cultured in 3F medium and subsequently grown in 3F medium or media containing FGF7 alone (‘differentiation media’) for 24 days. **(B)**Brightfield images of bud tip organoids growing in 3F media (left) or FGF7-differentiation media (right). Scale bar represents 500 μm. **(C)**NKX2.1 immunofluorescence of bud tip organoids grown in FGF7-differentiation media for 24 days. Scale bar represents 50 μm. **(D)** SOX9 and SOX2 immunofluorescence of bud tip organoids grown in FGF7-differentiation media for 24 days. Scale bar represents 50 μm. **(E)**QRT-PCR analysis of bud tip organoids grown in 3F or FGF7-differentiation media showing expression of several genes expressed in the lung epithelium. Data is shown as Fold Change relative to 3F-grown undifferentiated bud tip organoids. **(F)** QRT-PCR analysis of bud tip organoids grown in 3F or FGF7-differentiation media, along with whole distal fetal lung and cultured whole fetal intestine as reference tissues, showing expression of several genes expressed in the lung epithelium. Data is shown as Arbitrary Units. Values lower than 10^−3^ were considered undetected. **(F)** Immunostaining for airway markers in the adult human lung, and in bud tip organoids grown in FGF7-differentiation media. Markers are shown for goblet cells (MUC5AC, MUC5B), club cells (SCGB1A1, PLUNC) and neuroendocrine cells (SYN, CHGA). Scale bars represents 50 μm. **(G)** Transmission Electron Microscopy through a bud tip organoid grown in FGF7-differentiation media showing mucus filled vacuoles characteristic of goblet cells. Scale bar represents 100 nm. **(H)** Immunostaining for alveolar markers in the adult human lung, and in bud tip organoids grown in FGF7-differentiation media. Markers are shown for AECI cells (PDPN, HOPX) and AECII cells (Pro-SFTPC, SFTPB, ABCA3). Scale bars represent 50 μm. **(I)** Transmission Electron Microscopy of a human fetal lung at 13 weeks of gestation, and of a bud tip organoid grown in FGF7-differentiation media showing immature lamellar bodies surrounded by monoparticulate glycogen, characteristic of immature AT2 cells. Scale bars represent 100 nm. **(J-K)** Quantitation of cell type markers in bud tip organoids grow in differentiation media plotted as aggregate data ((H) numbers at top of bars represent positive cells/total cells counted across 5 individual organoids), and as individual data per organoid ((I) number of positive cells per organoid). Each data point in (I) represents an independent biological replicate and graphs indicate the mean +/− the standard error of the mean. **(L)** Summary of putative differentiated lung epithelial cell types generated *in vitro*.

**Figure 6:**
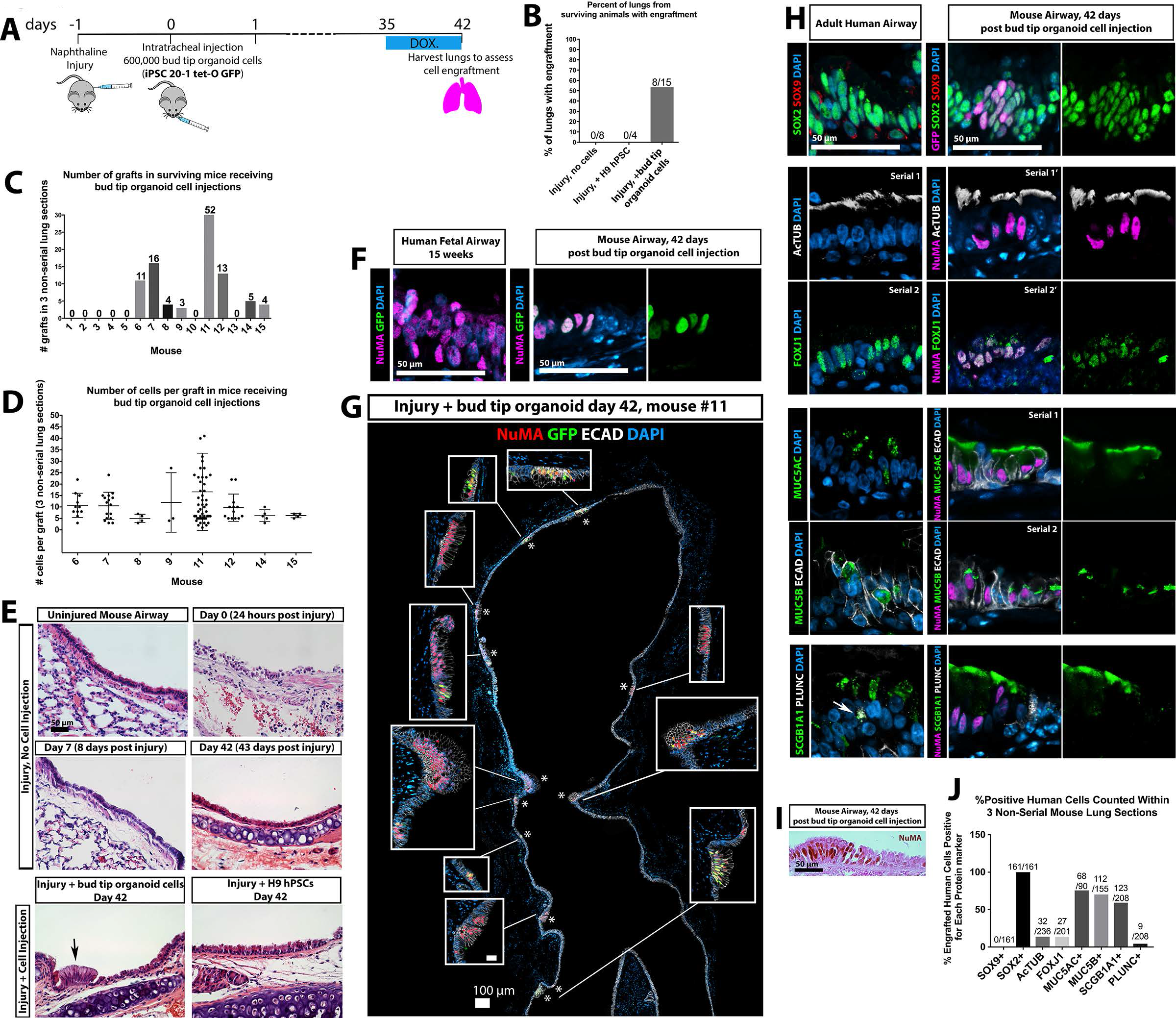
Engraftment of hPSC-derived bud tip progenitor organoid cells into the injured mouse airway. **(A)**Schematic of experimental design. Immunocompromised NSG male mice were injected with 300mg/kg of Naphthalene. 24 hours post-injury, mice were randomly assigned to receive an intratracheal injection of 600,000 single cells isolated from bud tip organoids generated from the iPSC 20-1 tet-O GFP line, undifferentiated H9 hPSCs, or no injection of cells. Doxycycline (1mg/ml) was added to the drinking water for the final week to induce expression of the tet-O GFP construct. Lungs were analyzed 6 weeks after cell injection. **(B)**Percent of lungs from surviving animals in each group exhibiting engraftment of human cells after 6 weeks, as determined by NuMA and GFP protein staining. **(C-D)**Engraftment was assessed based on human specific expression on NuMA and GFP in 3 independent histological sections from each surviving mouse. (C) The number of engraftment cell patches observed in 15 surviving animals in the group receiving bud tip organoid cells. (D) Quantitation of the number of human cells in each engrafted cell patch, in each mouse. Every data point represents the number of cells in a single patch of cells. **(E)**Hematoxylin & Eosin staining documenting the lung epithelial airway injury in control mice (injury, no cell injection), and in mice that received bud tip organoids, or undifferentiated iPSC injections. Engrafted patches of human cells were obvious (arrow), and were confirmed in serial sections using human specific antibodies. Scale bar for all images represents 50 μm. **(F)**Immunostaining for NuMA and GFP in human fetal lungs and in bud tip organoid transplanted lungs. Scale bar represents 50 μm. **(G)**Bud tip organoid transplanted lung showing several images stiched together to generate a large panel demonstrating multiple sites of engraftment, marked by NuMA and/or GFP, in the upper airway. High magnification insets are shown. Scale bar represents 100μm. **(H)**Immunostaining of adult human lung tissue and bud tip organoid transplanted lung tissue showing immunostaining for several lung epithelial markers, including SOX9 and SOX2, multiciliated cell markers Ac-TuB and FOXJ1, goblet cell markers MUC5AC and MUC5B, club cell markers SCGB1A1 and PLUNC. Scale bar represents 50 μm. **(I)** Bright field microscopic image showing immunohistochemistry for NuMA in a patch of engrafted cells, counterstained using eosin to visualize tufts of multiple cilia. Scale bar represents 50 μm. **(J)** Quantification of cell type markers in bud tip organoid transplanted lungs after 6 weeks. Data is plotted as aggregate data (numbers at top of bars represent positive cells/total cells counted across 3 engrafted lungs). Aggregated data from 3 non-serial sections for each mouse is plotted.

### *In vitro* differentiation

Since we were interested in investigating the potential of bud tip progenitor organoids to give rise to any lung epithelial lineage, we reasoned that withdrawing CHIR-99021 and RA may allow the cells to stochastically differentiate. Therefore, bud tip organoids were split into two treatment groups: 3F medium or FGF7-only for 23 days (Figure 5A). At the end of the experiment, control bud tip organoids maintained a clear, thin epithelium with a visible lumen, whereas FGF7-differentiated organoids appeared as denser cysts with mucous-like material inside of some organoids (Figure 5B). FGF7-differentiated organoids remained NKX2.1+/ECAD+ and SOX2+, signifying that they retained a lung epithelial identity, but showed decreased SOX9 protein expression (Figure 5C,D). Gene expression analysis using qRT-PCR to compare 3F (undifferentiated) vs. FGF7-differentiated organoids showed significant increases of genes expressed in differentiated airway and alveolar epithelium (Figure 5E). When further compared to the whole distal portion of human fetal lung (positive control) and cultured human fetal duodenum (negative control), differentiated bud tip organoids showed a significant increase relative to 3F in many genes associated with differentiated epithelial cells, with many genes trending towards levels seen in the whole fetal lung (Figure 5F).

Protein staining for airway and alveolar markers was carried out on human lung tissue samples as a reference and in FGF7-differentiated organoids (Figure 5G,I). We observed that cells within FGF7-differentiated organoids expressed proximal airway markers, including the goblet cell markers MUC5AC and MUC5B, the club cell markers SCGB1A1, and the neuroendocrine cell markers SNY and CHGA (Figure 5F). Of note, we observed that SCGB1A1 and PLUNC were co-expressed in only a small subset of club cells in the adult lung, whereas many club cells were marked only with SCGB1A1 (Figure 5G). We did not observe P63, FOXJ1 or cilia with Acetylated-Tubulin staining in FGF7-differentiated organoids (negative data not shown). Transmission electron microscopy (TEM) also revealed cells with mucous containing vesicles characteristic of goblet cells within FGF7-differentiated organoids (Figure 5G). Immunostaining for alveolar markers revealed cells positive for AECI markers PDPN and HOPX, as well as cells double positive for AECII markers Pro-SFTPC and SFTPB and Pro-SFTPC and ABCA3 (Figure 5I). Of note, TEM also revealed putative lamellar bodies that were similar to those seen in 13 week human fetal lungs (Figure 5J). In both tissues, putative lamellar bodies were characteristically surrounded by what appears to be monoparticulate glycogen (Stahlman et al., 2000). Both iPSC-derived and human fetal tissue lamellar bodies appear to be immature, and did not possess the typical concentric lamellae of mature structures (Vanhecke et al., 2010) (Figure 5J). Based on both IF and TEM data, it is most likely that the alveolar cell types observed in FGF7-differentiated organoids are reminiscent of an immature cell, and may require more specific conditions to further mature the primitive AECI and AECII cells.

Cells expressing cell type-specific lung epithelial markers were quantitated within FGF7-differentiated organoids, showing that organoids possessed cells expressing the alveolar markers: PDPN − 8.6% of total cells; HOPX − 5.9% of total cells; SFTPB ~45% of total cells; pro-SFTPC - ~ 45% of total cells; ABCA3 - 6.3% of total cells (Figure 5J-K). Many cells expressed airway markers: 43.2% of cells stained positive for nuclear SOX2, but not SOX9 (Figure 5J-K), 26.8% of cells stained positive for MUC5AC, while 7% of cells were positive for the club cell marker SCGB1A1, while FOXJ1 and P63 were absent (Figure 5K-L). Additionally, 2.0% of cells expressed very clear co-staining for Chromagranin A (CHGA) and Synaptophysin, markers for neuroendocrine cells (Figure 5G, K-L).

### In vivo engraftment

We next sought to explore the differentiation potential of hPSC-derived bud tip progenitor organoids by engrafting them into the airways of injured mouse lungs (Figure S6 and Figure 6). Recent studies have shown that damaging the lung epithelium promotes engraftment of adult human lung epithelial cells (Ghosh et al., 2017).

Interestingly, we observed that 7 days post injection of the cells, 79% of human cells in the mouse airway were still co-labeled by nuclear SOX2 and nuclear SOX9 and remained highly proliferative (Figure S6F-J). Immunofluorescent staining showed that of the small proportion of cells positive for only SOX2, 42.8% of engrafted human cells expressed the club cell marker SCGB1A1, and 25% expressed the goblet cell marker MUC5AC (Figure S6K-N). Multiciliated cells were not observed (negative FOXJ1 staining in Figure S6K) nor were basal cells nor alveolar cell markers (basal cells: P63, KRT5; AECI: HOPX, PDPN, RAGE; AECII: Pro-SFTPC, SFTPB, ABCA3; negative staining not shown). Collectively, our data suggested that hPSC-derived lung bud tip progenitor-like cells can engraft into the injured airway; however, the majority of engrafted cells (79%, Figure S6) retained expression of SOX9 after 7 days.

To determine whether engrafted cells could differentiate into multiple lung cell lineages, we carried out a long-term engraftment experiment where lungs were harvested 6 weeks after injection of cells (Figure 6). Mice received Naphthalene injury 24 hours prior to intratracheal injection of 600,000 dissociated bud tip organoid cells derived from iPSCs (iPSC20.1 tet-O GFP) (Figure 6A). Doxycycline was given on the final week of the experiment to induce expression from the tet-O-GFP transgene in the iPSC-derived cells. Experimental cohorts included: injury plus no cell injection (n=8 surviving animals); injury plus undifferentiated hPSC injection (n=4 surviving animals); injury plus bud tip organoid cell injection (n=15 surviving animals) (Figure 6B). Lungs from animals in all experimental groups successfully recovered from the injury (Figure 6E). Engrafted human cells were not observed in the “no cell injection” nor the “undifferentiated hPSC injection” cohorts (Figure 6B) as determined by NUMA+/GFP+ staining (Figure 6F-G). Of the 15 surviving mice that received bud tip organoid injections, 8 of them showed patches of NUMA+/GFP+ cells that engrafted into the airway (Figure 6C-D, 6F-G). The number of engrafted cell patches (Figure 6C) and the number of cells per patch (Figure 6D) varied across individual mice. While 2 mice exhibited a total of 4 small patches of engrafted cells in the bronchioles, the overwhelming majority of engrafted cells were found in the trachea and primary/secondary bronchi of mice. The rare engrafted cells within the bronchioles stained positive for proximal cell markers SOX2, MUC5AC or FOXJ1/AcTUB (data not shown).

We conducted immunostaining for several lung epithelial markers to determine if engrafted tissue had differentiated, and we compared staining patterns to adult human lung tissue (Figure 6H). Immunostaining of the injury group that did not receive cells was also examined (Figure S6Q). We noted that engrafted human cells expressed low levels of NKX2.1, consistent with the human airway where the majority of cells expressed nearly undetectable levels of NKX2.1 (Figure S6S-T). Engrafted human cells (as determined by GFP or NuMA expression) also expressed SOX2, but not SOX9, suggesting these cells had differentiated into an airway fate (Figure 6H). Consistent with this observation, we observed engrafted human cells that possessed multiple cilia and co-expressed NuMA and multiciliated cell markers, AcTub and FOXJ1 (Figure 6H) and as determined in histological sections (Figure 6I). We also observed human cells expressing goblet cell markers MUC5AC and MUC5B. Finally, we observed human cells expressing the club cell marker SCGB1A1, but not PLUNC. Interestingly, we noted that PLUNC only marked a small subset of club cells in the human airway, suggesting that there is an underappreciated heterogeneity within this population in the human lung (Figure 6H). We did not find evidence that engrafted human cells expressed the basal cell marker P63 nor alveolar cell specific markers (negative data not shown).

## Discussion

The ability to study human lung development, homeostasis and disease is limited by our ability to carry out functional experiments in human tissues. This has led to the development of many different *in vitro* model systems using primary human tissue, and using cells and tissues derived from hPSCs (Dye et al., 2016b; Miller and Spence, 2017). Current approaches to differentiate hPSCs have used many techniques, including the stochastic differentiation of lung-specified cultures into many different lung cell lineages (Chen et al., 2017; Huang et al., 2013; Wong et al., 2012), FACS based sorting methods to purify lung progenitors from mixed cultures followed by subsequent differentiation (Gotoh et al., 2014; Konishi et al., 2015; Longmire et al., 2012; McCauley et al., 2017), and by expanding 3-dimensional ventral foregut spheroids into lung organoids (Dye et al., 2016a; Dye 2015). Many or all, of these approaches rely on the differentiation of a primitive NKX2.1+ lung progenitor cell population, followed by subsequent differentiation into different cell lineages. However, during normal development, early NKX2.1-specified lung progenitors transition through a SOX9+ bud tip progenitor cell state on their way to terminal differentiation into both alveolar and airway cell fates (Branchfield et al., 2015; Rawlins et al., 2009). Despite the developmental importance of this bud tip progenitor, differentiation of a similar progenitor from hPSCs has remained elusive. For example, several studies have identified robust methods to sort and purify lung epithelial progenitor cells from a mixed population (Gotoh et al., 2014; Konishi et al., 2015; McCauley et al., 2017); however, whether or not these populations represent or transition through an epithelial bud tip-like state is unknown.

Inducing and maintaining the bud tip progenitor state in hPSC-derivatives *in vitro* has remained elusive, in part, because the complex signaling networks required controlling this population are not well understood. A recent study demonstrated the ability to culture human fetal bud tips *in vitro*, supporting the findings of the current study (Nikolić et al., 2017), however *bona fide* SOX9+/SOX2+ bud tip progenitor-like cells derived from hPSCs have not been described nor well characterized prior to this work. The ability to induce, *de novo*, robust populations of cells from hPSCs that do not rely on specialized sorting or purification protocols suggest that biologically robust experimental findings can be used in a manner that predicts how a naïve cell will behave in the tissue culture dish with a high degree of accuracy, and across multiple cell lines (D'Amour et al., 2005; Green et al., 2011; Spence et al., 2011).

Our studies also identified significant species-specific differences between the human and fetal mouse lung. Differences included both gene/protein expression differences, as well as functional differences when comparing how cells responded to diverse signaling environments *in vitro*. These mouse-human differences highlight the importance of validating observations made in hPSC-derived tissues by making direct comparisons with human tissues, as predictions or conclusions about human cell behavior based on results generated from the developing mouse lung may be misleading.

Our experimental findings, in combination with previously published work, have also raised new questions that may point to interesting avenues for future mechanistic studies to determine how specific cell types of the lung are generated. Previously, we have shown that lung organoids grown in high concentrations of FGF10 predominantly give rise to airway-like tissues, with a small number of alveolar cell types and a lack of alveolar structure, and these organoids also possess abundant mesenchymal cell populations (Dye et al., 2016a; Dye 2015). Here, our results suggest that high concentrations of FGF10 alone do not play a major role in supporting robust growth of epithelial bud tip progenitor cells. We also note that lung organoids grown in high FGF10 possess abundant P63+ basal-like cells (Dye et al., 2015), whereas bud tip organoids grown in 3F medium lack this population. These findings suggest that we still do not fully appreciate how various signaling pathways interact to control cell fate decisions or expansion of mesenchymal populations, and lay the groundwork for many future studies.

Taken together, our current work has identified a signaling network required for the induction, expansion and maintenance of hPSC-derived lung epithelial bud tip progenitors. Simple needle passaging allowed us to expand a nearly homogenous population of proliferative bud tip-like progenitor cells for over 16 weeks in culture, which remained multipotent *in vitro* and which were able to engraft into injured mouse lungs, terminally differentiate, and respond to systemic factors. The current thus study offers a robust and reproducible method to generate and maintain epithelial bud tip progenitors, which will facilitate future studies aimed at elucidating fundamental developmental mechanisms regulating human lung progenitor cells, and which may have applicability to regenerative medicine in the future.

## Methods

### EXPERIMENTAL MODEL AND SUBJECT DETAILS

#### Mouse models

All animal research was approved by the University of Michigan Committee on Use and Care of Animals. Lungs from Sox9-eGFP (MGI ID:3844824), Sox9CreER;Rosa^Tomato/Tomato^ (MGI ID:5009223 and 3809523)(Kopp et al., 2011), or wild type lungs from CD1 embryos (Charles River) were dissected at embryonic day (E) 13.5, and buds were isolated as described below and as previously described (del Moral and Warburton, 2010). 8-10 week old Immunocompromised NSG mice (Jackson laboratories strain #0005557) were used for engraftment studies. Pilot studies identified that females were more sensitive to Naphthalene and died at a higher rate, therefore male mice were used for engraftment experiments.

#### Human lung tissue

All research utilizing human fetal tissue was approved by the University of Michigan institutional review board. Normal human fetal lungs were obtained from the University of Washington Laboratory of Developmental Biology, and epithelial bud tips were dissected as described below. All tissues were shipped overnight in Belzer’s solution at 4 degrees Celsius and were processed and cultured within 4 hours of obtaining the specimen. Experiments to evaluate the effect on progenitor maintenance in culture by media conditions were repeated using tissues from 3 individual lung specimens; (1) 84 day post fertilization of unknown gender, (2), 87 day post fertilization male, and (3), 87 day post fertilization of unknown gender. RNAseq experiments of whole lung homogenate utilized tissue from 3 additional individual lungs; (4) 59 day male (5) 87 day of unknown gender and (6) 125 day of unknown gender. Use of normal adult human lung tissue was approved by University of Michigan institutional review board, and was obtained from warm autopsy organ donors arranged by the Gift of Life, Michigan. Adult human RNA-sequencing samples representing bulk sequencing of whole lung homogenates were obtained from the EMBL-EBI ArrayExpress repository (https://www.ebi.ac.uk/arrayexpress/experiments/E-MTAB-1733/) (Fagerberg et al., 2014).

#### Cell lines and culture conditions

##### Mouse and human primary cultures

Isolated mouse bud tips were cultured in 4-6 μl droplets of matrigel, covered with media, and kept at 37 degrees Celsius with 5% CO2. Isolated human fetal lung bud tips were cultured in 25-50 μl droplets of matrigel, covered with media, and kept at 37 degrees Celsius with 5% CO2. Cultures media was changed every 2-4 days.

##### Generation and culture of hPSC-derived lung organoids

The University of Michigan Human Pluripotent Stem Cell Research Oversight (hPSCRO) Committee approved all experiments using human embryonic and induced pluripotent stem cell (hESC, iPSC) lines. Patterned lung organoids were generated from 4 independent pluripotent cell lines in this study: hESC line UM63-1 (NIH registry #0277) was obtained from the University of Michigan and hESC lines H9 and H1 (NIH registry #0062 and #0043, respectively) were obtained from the WiCell Research Institute. iPSC20.1 was previously described (Spence et al., 2011). ES cell lines were routinely karyotyped to ensure normal karyotype and ensure the sex of each line (H9 - XX, UM63-1 - XX, H1 - XY). All cell lines are routinely monitored for mycoplasma infection monthly using the MycoAlert Mycoplasma Detection Kit (Lonza). Stem cells were maintained on hESC-qualified Matrigel (Corning Cat# 354277) using mTesR1 medium (Stem Cell Technologies). hESCs were maintained and passaged as previously described (Spence et al., 2011) and ventral foregut spheroids were generated as previously described (Dye et al., 2016a; Dye 2015). Following differentiation, free-floating foregut spheroids were collected from differentiated stem cell cultures and plated in a matrigel droplet on a 24-well tissue culture grade plate.

A summary of different tissue/sample types, nomenclature and descriptions can be found in Table 1.

### METHOD DETAILS

#### Isolation and culture of mouse lung epithelial buds

Mouse buds were dissected from E13.5 embryos. For experiments using Sox9CreER;Rosa^Tomato/Tomato^ mice, 50 ug/g of tamoxifen was dissolved in corn oil and given by oral gavage on E12.5, 24 hours prior to dissection. Briefly, in a sterile environment, whole lungs were placed in 100% dispase (Corning Cat# 354235) on ice for 30 minutes. Lungs were then transferred using a 25uL wiretrol tool (Drummond Scientific Cat# 5-000-2050) to 100% FBS (Corning Cat#35-010-CV) on ice for 15 minutes, and then transferred to a solution of Dulbecco’s Modified Eagle Medium: Nutrient Mixture F12 (DMEM/F12, ThermoFisher SKU# 12634-010) with 10% FBS and 1x Pennicillin-Streptomycin (ThermoFisher Cat# 15140122) on ice. To dissect buds, a single lung or lung lobe was transferred by wiretrol within a droplet of media to a 100mm sterile petri dish. Under a dissecting microscope, the mesenchyme was carefully removed and epithelial bud tips were torn away from the bronchial tree using tungsten needles (Point Technologies, Inc.). Care was taken to remove the trachea and any connective tissue from dissected lungs. Isolated bud tips were picked up using a p20 pipette and added to an eppendorf tube with cold Matrigel (Corning Ref# 354248) on ice. The buds were picked up in a p20 pipette with 4-6 uL of Matrigel and plated on a 24-well tissue culture well (ThermoFisher Cat# 142475). The plate was moved to a tissue culture incubator and incubated for 5 minutes at 37 degrees Celsius and 5% CO2 to allow the Matrigel droplet to solidify. 500uL of media was then added to the dish in a laminar flow hood. Media was changed every 2-3 days.

#### Isolation and culture of human fetal lung epithelial bud tips

Distal regions of 12 week fetal lungs were cut into ~2mm sections and incubated with dispase, 100% FBS and then 10% FBS as described above, and moved to a sterile petri dish. Mesenchyme was removed by repeated pipetting of distal lung pieces after dispase treatment. Buds were washed with DMEM until mesenchymal cells were no longer visible. Buds were then moved to a 1.5 mL eppendorf tube containing 200 μL of Matrigel, mixed using a p200 pipette, and plated in ~20 μL droplets in a 24 well tissue culture plate. Plates were placed at 37 degrees Celsius with 5% CO2 for 20 minutes while droplets solidified. 500uL of media was added to each well containing a droplet. Media was changed every 2-4 days.

#### RNA-sequencing and Bioinformatic Analysis

RNA was isolated using the mirVana RNA isolation kit, following the “Total RNA” isolation protocol (Thermo-Fisher Scientific, Waltham MA). RNA sequencing library preparation and sequencing was carried out by the University of Michigan DNA Sequencing Core and Genomic Analysis Services (https://seqcore.brcf.med.umich.edu/). 50bp single end cDNA libraries were prepared using the Truseq RNA Sample Prep Kit v2 (Illumina), and samples were sequenced on an Illumina HiSeq 2500. Transcriptional quantitation analysis was conducted using 64-bit Debian Linux stable version 7.10 (“Wheezy”). Pseudoalignment of RNA-sequencing reads was computed using kallisto v0.43.0 and a normalized data matrix of pseudoaligned sequences (Transcripts Per Million, TPM) and differential expression was calculated using the R package DEseq2 (Bray et al., 2016; Love et al., 2014). Data analysis was performed using the R statistical programming language (http://www.R-project.org/) and was carried out as previously described (Dye et al., 2015; Finkbeiner et al., 2015; Tsai et al., 2016). The complete sequence alignment, expression analysis and all corresponding scripts can be found at https://github.com/hilldr/Miller_Lung_Organoids_2017. All raw data files generated by RNA-sequencing have been deposited to the EMBL-EBI ArrayExpress database (accession ID: E-MTAB-6023).

#### Naphthaline injury

Naphthaline (Sigma #147141) was dissolved in corn oil at a concentration of 40 mg/ml. Adult male NSG mice were chosen for these experiments because we observed improved recovery and survival following injury compared to female mice. Mice that were 8-10 weeks of age were given i.p. injections at a dose of 300 mg/kg weight.

#### Intratracheal injection of fetal progenitor organoids and hPSC-derived bud tip organoid cells into immunocompromised mouse lungs

##### Generating single cells from organoid tissues

2-3 matrigel droplets containing organoid tissues were removed from the culture plate and combined in a 1.5mL eppendorf tube with 1mL of room temperature Accutase (Sigma #A6964). The tube was laid on its side to prevent organoids from settling to the bottom. Tissue was pipetted up and down 15-20 times with a 1mL tip every 5 minutes for a total of 20 minutes. Single cell status was determined by microscopic observation using a hemocytometer. Cells were diluted to a concentration of 500,000-600,000 cells per 0.03 mL in sterile PBS.

##### Intratracheal injection of cells

Injection of cells into the mouse trachea was performed as previously described (Badri et al., 2011; Cao et al., 2017). Briefly, animals were anesthetized and intubated. Animals were given 500,000-600,000 single cells in 30-35 μL of sterile PBS through the intubation cannula.

#### Culture media, growth factors and small molecules

##### Serum-free basal media

All mouse bud, human fetal bud, and hPSC-derived human lung organoids were grown in serum-free basal media (basal media) with added growth factors. Basal media consisted of Dulbecco’s Modified Eagle Medium: Nutrient Mixture F12 (DMEM/F12, ThermoFisher SKU# 12634-010) supplemented with 1X N2 supplement (ThermoFisher Catalog# 17502048) and 1X B27 supplement (ThermoFisher Catalog# 17504044), along with 2mM Glutamax (ThermoFisher Catalog# 35050061), 1x Pennicillin-Streptomycin (ThermoFisher Cat# 15140122) and 0.05% Bovine Serum Albumin (BSA; Sigma product# A9647). BSA was weighed and dissolved in DMEM F/12 media before being passed through a SteriFlip 0.22 uM filter (Millipore Item# EW-29969-24) and being added to basal media. Media was stored at 4 degrees Celsius for up to 1 month. On the day of use, basal media was aliquoted and 50ug/mL Ascorbic acid and 0.4 uM Monothioglycerol was added. Once ascorbic acid and monothioglycerol had been added, media was used within one week.

##### Growth factors and small molecules

Recombinant Human Fibroblast Growth Factor 7 (FGF7) was obtained from R&D systems (R&D #251-KG/CF) and used at a concentration of 10 ng/mL unless otherwise noted. Recombinant Human Fibroblast Growth Factor 10 (FGF10) was obtained either from R&D systems (R&D #345-FG) or generated in house (see below), and used at a concentration of 10 ng/mL (low) or 500 ng/mL (high) unless otherwise noted. Recombinant Human Bone Morphogenic Protein 4 (BMP4) was purchased from R&D systems (R&D Catalog # 314-BP) and used at a concentration of 10 ng/mL. All-trans Retinoic Acid (RA) was obtained from Stemgent (Stemgent Catalog# 04-0021) and used at a concentration of 50 nM. CHIR-99021, a GSK3β inhibitor that stabilizes β-CATENIN, was obtained from STEM CELL technologies (STEM CELL Technologies Catalog# 72054) and used at a concentration of 3 uM. Y27632, a ROCK inhibitor (APExBIO Cat# A3008) was used at a concentration of 10uM.

##### Generation and Isolation of human recombinant FGF10

Recombinant human FGF10 was produced in-house. The recombinant human FGF10 (rhFGF10) expression plasmid pET21d-FGF10 in *E. coli* strain BL21*trx*B(DE3) was a gift from James A. Bassuk at the University of Washington School of Medicine (Bagai et al., 2002). *E. coli* was grown in standard LB media with peptone derived from meat, carbenicillin and glucose. rhFGF10 expression was induced by addition of isopropyl-1-thio-β-D-galactopyranoside (IPTG). rhFGF10 was purified by using a HiTrap-Heparin HP column (GE Healthcare, 17040601) with step gradients of 0.5M to 2M LiCl. From a 200 ml culture, 3 mg of at least 98% pure rFGF-10 (evaluated by SDS-PAGE stained with Coomassie Blue R-250) was purified. rFGF10 was compared to commercially purchased human FGF10 (R&D Systems) to test/validate activity based on the ability to phosphorylate ERK1/2 in an A549 alveolar epithelial cell line (ATCC Cat#CCL-185) as assessed by western blot analysis.

#### RNA extraction and quantitative RT-PCR analysis

RNA was extracted using the MagMAX-96 Total RNA Isolation System (Life Technologies). RNA quality and concentration was determined on a Nanodrop 2000 spectrophotometer (Thermo Scientific). 100 ng of RNA was used to generate a cDNA library using the SuperScript VILO cDNA master mix kit (Invitrogen) according to manufacturer’s instructions. qRT-PCR analysis was conducted using Quantitect SYBR Green Master Mix (Qiagen) on a Step One Plus Real-Time PCR system (Life Technologies). Expression was calculated as a change relative to GAPDH expression using arbitrary units, which were calculated by the following equation: [2^(GAPDH Ct - Gene Ct)] × 10,000. A Ct value of 40 or greater was considered not detectable. A list of primer sequences used can be found in Table 2.

**Table 2:**
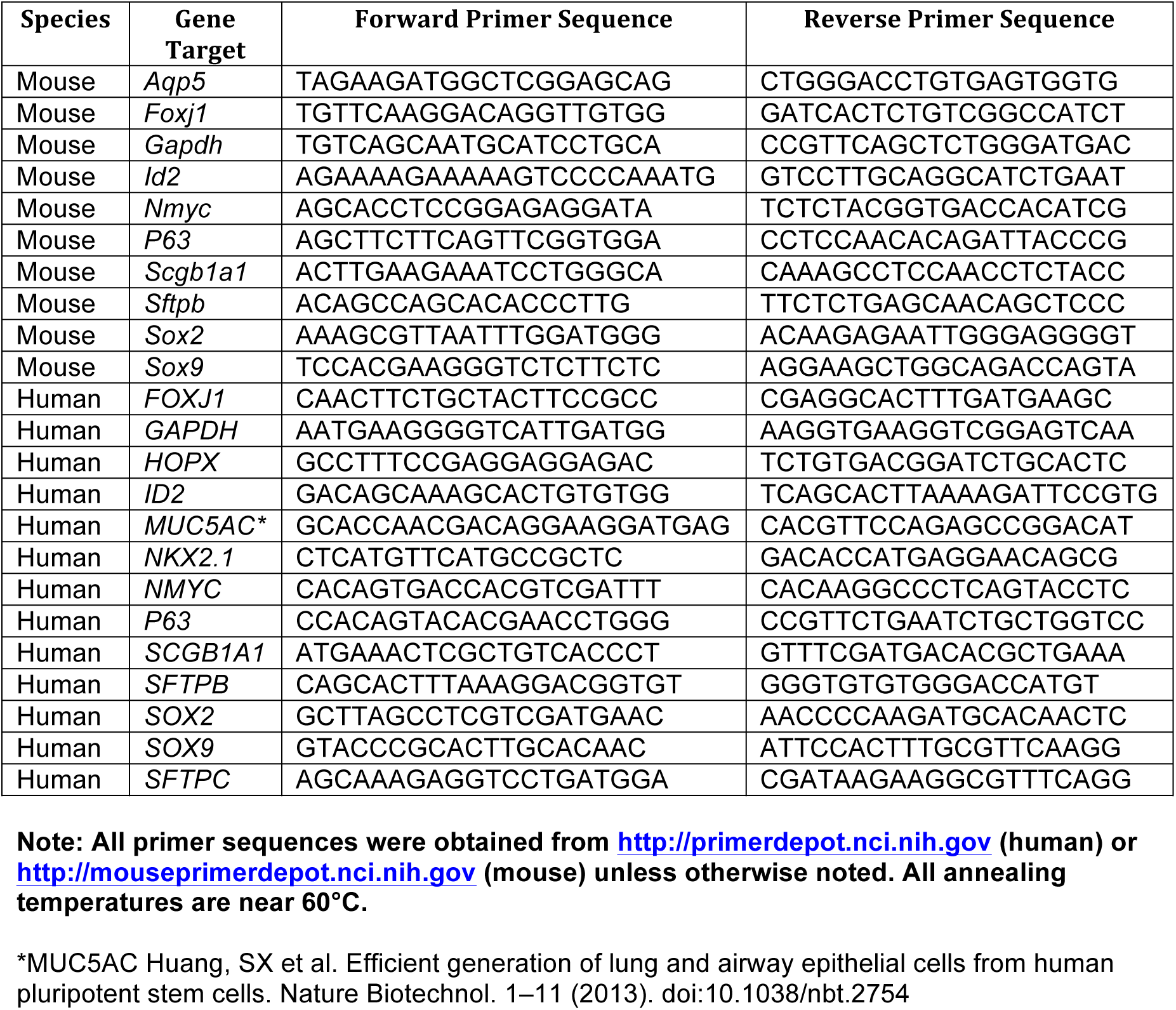
Qrt-PCR primer sequences.

#### Tissue preparation, Immunohistochemistry, Electron Microscopy and imaging

##### Paraffin sectioning and staining

Mouse bud, human bud, and HLO tissue was fixed in 4% Paraformaldehyde (Sigma) for 2 hours and rinsed in PBS overnight. Tissue was dehydrated in an alcohol series, with 30 minutes each in 25%, 50%, 75% Methanol:PBS/0.05% Tween-20, followed by 100% Methanol, and then 100% Ethanol. Tissue was processed into paraffin using an automated tissue processor (Leica ASP300). Paraffin blocks were sectioned 5-7 uM thick, and immunohistochemical staining was performed as previously described (Spence et al., 2009). A list of antibodies, antibody information and concentrations used can be found in Table 3. PAS Alcian blue staining was performed using the Newcomer supply Alcian Blue/PAS Stain kit (Newcomer Supply, Inc.) according to manufacturer’s instructions.

**Table 3:**
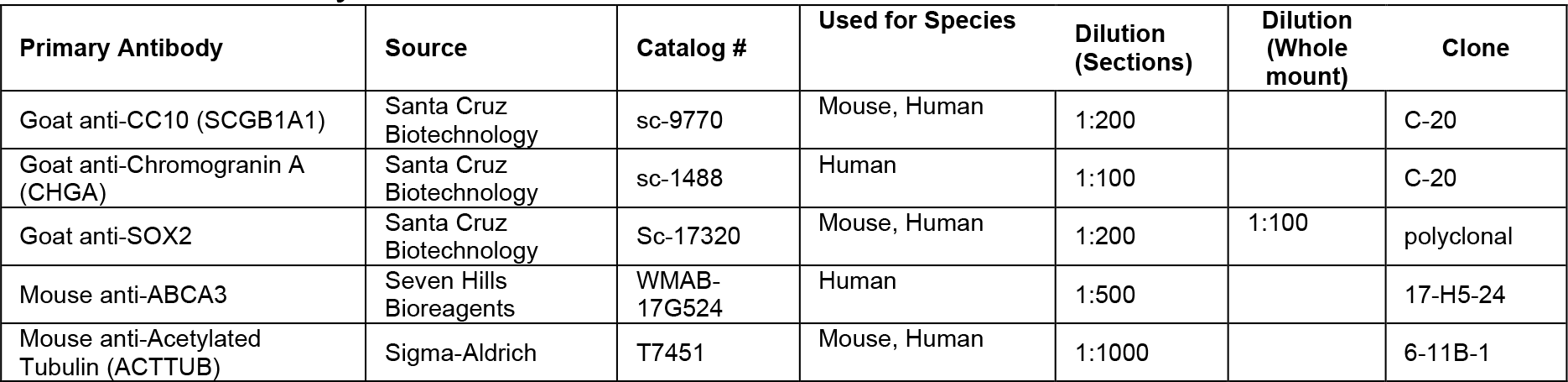
Antibody information.

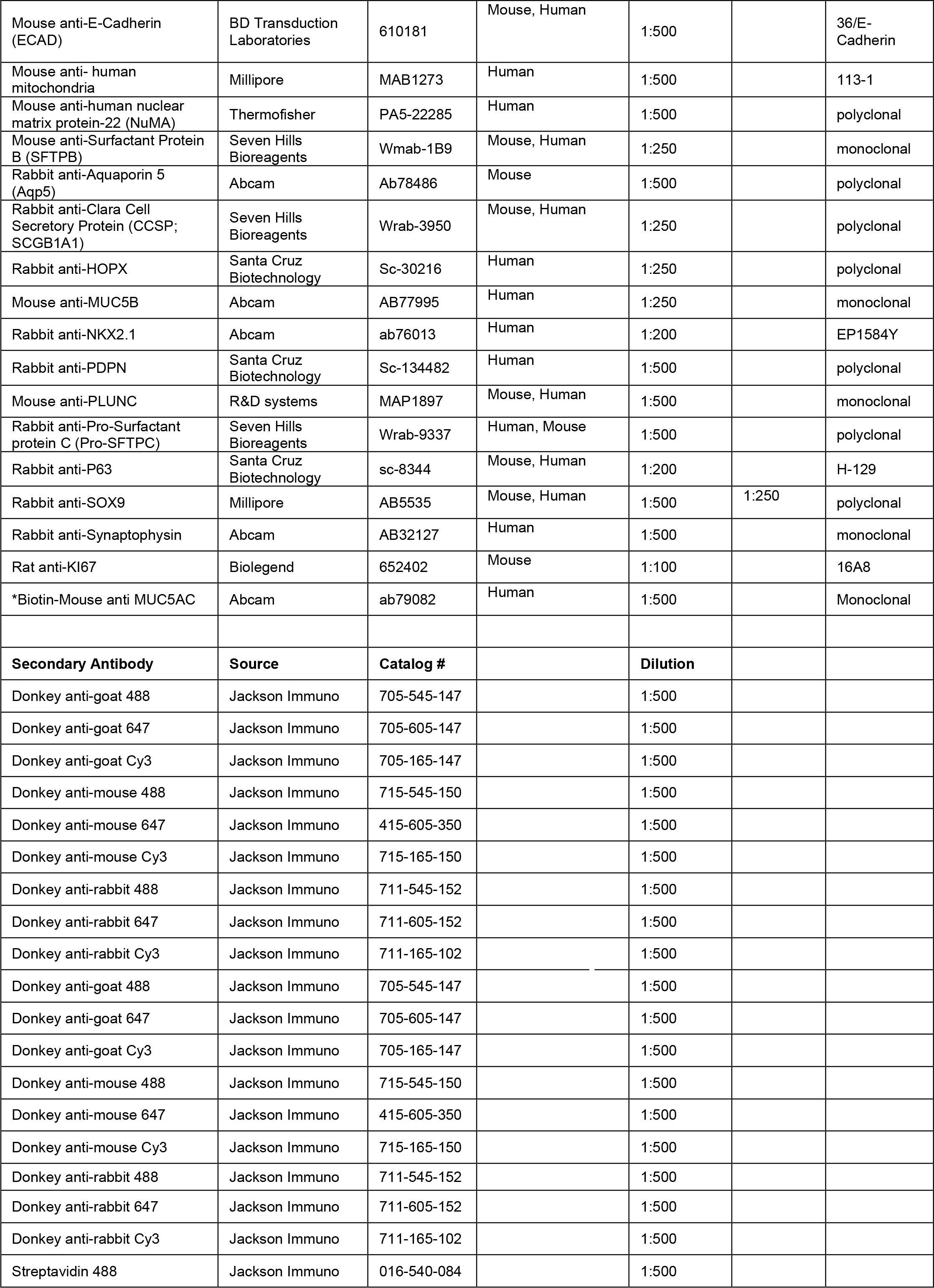

##### Whole mount staining

For whole mount staining tissue was placed in a 1.5mL eppendorf tube and fixed in 4% paraformaldehyde (Sigma) for 30 minutes. Tissue was then washed with PBS/0.05% Tween-20 (Sigma) for 5 hours, followed by a 2.5-hour incubation with blocking serum (PBS-Tween-20 plus 5% normal donkey serum). Primary antibodies were added to blocking serum and tissue was incubated for at least 24 hours at 4 degrees Celcius. Tissue was then washed for 5 hours with several changes of fresh PBS-Tween-20. Secondary antibodies were added to fresh blocking solution and tissue was incubated for 12-24 hours, followed by 5 hours of PBS-Tween-20 washes. Tissue was then dehydrated to 100% methanol and carefully moved to the center of a single-well EISCO concave microscope slide (ThermoFisher Cat# S99368) using a glass transfer pipette. 5-7 drops of Murray’s clear (2 parts Benzyl alcohol, 1 part Benzyl benzoate [Sigma]) were added to the center of the slide, and slides were coverslipped and sealed with clear nail polish.

##### In situ hybridization

*In situ* hybridization for ID2 was performed using the RNAscope 2.5 HD manual assay with brown chromogenic detection (Advanced Cell Diagnostics, Inc.) according to manufacturers instructions. The human 20 base pair ID2 probe was generated by Advanced Cell Diagnostics targeting 121-1301 of ID2 (gene accession NM_002166.4) and is commercially available.

##### Imaging and image processing

Images of fluorescently stained slides were taken on a Nikon A-1 confocal microscope. When comparing groups within a single experiment, exposure times and laser power were kept consistent across all images. All Z-stack imaging was done on a Nikon A-1 confocal microscope and Z-stacks were 3-D rendered using Imaris software. Brightness and contrast adjustments were carried out using Adobe Photoshop Creative Suite 6 and adjustments were made uniformly across images. Low magnification brightfield images of live cultures were taken using an Olympus S2X16 dissecting microscope. Image brightness and contrast was enhanced equally for all images within a single experiment using Adobe Photoshop. Images were cropped where noted in figure legends to remove blank space surrounding buds or cultures.

##### Transmission Electron Microscopy

Undifferentiated and differentiated iPSC-derived bud tip organoids and 13 week human fetal lung tissue was fixed overnight in 2.5% glutaraldehyde in Sorensen’s buffer (0.1M). The following day tissue was washed with Cacodylate buffer (0.05M) then post fixed for one hour in 1% osmium tetroxide. After fixation, the tissue was rinsed again in Cacodylate buffer (0.05M). The tissue was then dehydrated in an ethanol gradient, with tissue incubating for 15 minutes in solutions of 25%, 50%, 75% and 100% ethanol. Tissue was then cleared in propylene oxide. Epon resin was infiltrated into the tissue by mixing propylene oxide and Epon resin (3:1, 1:1, 1:3), tissues were incubated in each mixture for 24 hours on a rocker. Tissues were then submerged into full strength Epon resin and rocked for two more days at room temperature, changing to fresh resin each day. Using fresh Epon resin, the tissues were arranged in molds and allowed to polymerize for 24 hours in a 60°C oven. Sections were cut using a Leica EM UC7 ultramicrotome and imaged using a JEOL 1400-Plus Transmission Electron Microscope.

#### Quantification and Statistical Analysis

All plots and statistical analysis were done using Prism 6 Software (GraphPad Software, Inc.). For statistical analysis of qRT-PCR results, at least 3 biological replicates for each experimental group were analyzed and plotted with the standard error of the mean. If only two groups were being compared, a two-sided student’s T-test was performed. In assessing the effect of length of culture with FGF7 on gene expression in mouse buds (Figure S2G), a one-way, unpaired Analysis of Variance (ANOVA) was performed for each individual gene over time. The mean of each time point was compared to the mean of the expression level for that gene at day 0 of culture. If more than two groups were being compared within a single experiment, an unpaired one-way analysis of variance was performed followed by Tukey’s multiple comparison test to compare the mean of each group to the mean of every other group within the experiment. For all statistical tests, a significance value of 0.05 was used. For every analysis, the strength of p values is reported in the figures according the following: P > 0.05 ns, P ≤ 0.05 *, P ≤ 0.01 **, P ≤ 0.001 ***, P ≤ 0.0001 ****. Details of statistical tests can be found in the figure legends.

## Acknowledgements

JRS is supported by the NIH-NHLBI (R01 HL119215). AJM is supported by the NIH Cellular and Molecular Biology training grant at Michigan (T32 GM007315), and by the Tissue Engineering and Regeneration Training Grant (DE00007057-40). The University of Washington Laboratory of Developmental Biology was supported by NIH Award Number 5R24HD000836 from the Eunice Kennedy Shriver National Institute of Child Health & Human Development.

## Notes

Conflicts of Interest: The authors have no conflicts to declare.

## References

Abler, L. L., Mansour, S. L., Sun, X., 2009. Conditional gene inactivation reveals roles for Fgf10 and Fgfr2 in establishing a normal pattern of epithelial branching in the mouse lung. Dev. Dyn. 238, 1999–2013. doi:10.1002/dvdy.22032

Badri, L., Walker, N. M., Ohtsuka, T., Wang, Z., Delmar, M., Flint, A., Peters-Golden, M., Toews, G. B., Pinsky, D. J., Krebsbach, P. H., Lama, V. N., 2011. Epithelial Interactions and Local Engraftment of Lung-Resident Mesenchymal Stem Cells. American Journal of Respiratory Cell and Molecular Biology 45, 809–816. doi:10.1165/rcmb.2010-0446OC

Bagai, S., Rubio, E., Cheng, J.-F., Sweet, R., Thomas, R., Fuchs, E., Grady, R., Mitchell, M., Bassuk, J. A., 2002. Fibroblast growth factor-10 is a mitogen for urothelial cells. J. Biol. Chem. 277, 23828–23837. doi: 10.1074/jbc.M201658200

Bellusci, S., Furuta, Y., Rush, M. G., Henderson, R., Winnier, G., Hogan, B. L., 1997a. Involvement of Sonic hedgehog (Shh) in mouse embryonic lung growth and morphogenesis. Development 124, 53–63.

Bellusci, S., Grindley, J., Emoto, H., Itoh, N., Hogan, B. L., 1997b. Fibroblast growth factor 10 (FGF10) and branching morphogenesis in the embryonic mouse lung. Development 124, 4867–4878.

Bellusci, S., Henderson, R., Winnier, G., Oikawa, T., Hogan, B. L., 1996. Evidence from normal expression and targeted misexpression that bone morphogenetic protein (Bmp-4) plays a role in mouse embryonic lung morphogenesis. Development 122, 1693–1702.

Branchfield, K., Li, R., Lungova, V., Verheyden, J. M., McCulley, D., Sun, X., 2015. A three-dimensional study of alveologenesis in mouse lung. Developmental Biology. doi: 10.1016/j.ydbio.2015.11.017

Bray, N. L., Pimentel, H., Melsted, P., Pachter, L., 2016. Near-optimal probabilistic RNA-seq quantification. Nat Biotechnol 34, 525–527. doi:10.1038/nbt.3519

Cao, P., Aoki, Y., Badri, L., Walker, N. M., Manning, C. M., Lagstein, A., Fearon, E. R., Lama, V. N., 2017. Autocrine lysophosphatidic acid signaling activates β-catenin and promotes lung allograft fibrosis. J. Clin. Invest. 127, 1517–1530. doi:10.1172/JCI88896

Chang, D. R., Martinez Alanis, D., Miller, R. K., Ji, H., Akiyama, H., McCrea, P. D., Chen, J., 2013. Lung epithelial branching program antagonizes alveolar differentiation. Proceedings of the National Academy of Sciences. doi: 10.1073/pnas.1311760110

Chen, Y.-W., Huang, S. X., de Carvalho, A. L.R.T., Ho, S.-H., Islam, M. N., Volpi, S., Notarangelo, L. D., Ciancanelli, M., Casanova, J.-L., Bhattacharya, J., Liang, A. F., Palermo, L. M., Porotto, M., Moscona, A., Snoeck, H.-W., 2017. A three-dimensional model of human lung development and disease from pluripotent stem cells. Nat. Cell Biol. 19, 542–549. doi:10.1038/ncb3510

Cornett, B., Snowball, J., Varisco, B. M., Lang, R., Whitsett, J., Sinner, D., 2013. Wntless is required for peripheral lung differentiation and pulmonary vascular development. Developmental Biology 379, 38–52. doi:10.1016/j.ydbio.2013.03.010

D’Amour, K. A., Agulnick, A. D., Eliazer, S., Kelly, O. G., Kroon, E., Baetge, E. E., 2005. Efficient differentiation of human embryonic stem cells to definitive endoderm. Nat Biotechnol 23, 1534–1541. doi:10.1038/nbt1163

Danopoulos, S., Alonso, I., Thornton, M., Grubbs, B., Bellusci, S., Warburton, D., Alam, Al, D., 2017. Human lung branching morphogenesis is orchestrated by the spatio-temporal distribution of ACTA2, SOX2 and SOX9. Am. J. Physiol. Lung Cell Mol. Physiol. ajplung.00379.2017-17. doi:10.1152/ajplung.00379.2017

del Moral, P.-M., Warburton, D., 2010. Explant culture of mouse embryonic whole lung, isolated epithelium, or mesenchyme under chemically defined conditions as a system to evaluate the molecular mechanism of branching morphogenesis and cellular differentiation. Methods Mol. Biol. 633, 71–79. doi:10.1007/978-1-59745-019-5_5

Desai, T. J., Chen, F., Lü, J., Qian, J., Niederreither, K., Dollé, P., Chambon, P., Cardoso, W. V., 2006. Distinct roles for retinoic acid receptors alpha and beta in early lung morphogenesis. Developmental Biology 291, 12–24. doi:10.1016/j.ydbio.2005.10.045

Desai, T. J., Malpel, S., Flentke, G. R., Smith, S. M., Cardoso, W. V., 2004. Retinoic acid selectively regulates Fgf10 expression and maintains cell identity in the prospective lung field of the developing foregut. Developmental Biology 273, 402–415. doi:10.1016/j.ydbio.2004.04.039

Domyan, E. T., Ferretti, E., Throckmorton, K., Mishina, Y., Nicolis, S. K., Sun, X., 2011. Signaling through BMP receptors promotes respiratory identity in the foregut via repression of Sox2. Development 138, 971–981. doi:10.1242/dev.053694

Domyan, E. T., Sun, X., 2010. Patterning and plasticity in development of the respiratory lineage. Dev. Dyn. 240, 477–485. doi:10.1002/dvdy.22504

Dye, B. R., Dedhia, P. H., Miller, A. J., Nagy, M. S., White, E. S., Shea, L. D., Spence, J. R., Rossant, J., 2016a. A bioengineered niche promotes in vivo engraftment and maturation of pluripotent stem cell derived human lung organoids. Elife 5, e19732. doi:10.7554/eLife.19732

Dye, B. R., Hill, D. R., Ferguson, M. A., Tsai, Y.-H., Nagy, M. S., Dyal, R., Wells, J. M., Mayhew, C. N., Nattiv, R., Klein, O. D., White, E. S., Deutsch, G. H., Spence, J. R., 2015. In vitro generation of human pluripotent stem cell derived lung organoids. Elife 4. doi:10.7554/eLife.05098

Dye, B. R., Miller, A. J., Spence, J. R., 2016b. How to Grow a Lung: Applying Principles of Developmental Biology to Generate Lung Lineages from Human Pluripotent Stem Cells. Curr Pathobiol Rep 4, 1–11. doi:10.1007/s40139-016-0102-x

Fagerberg, L., Hallström, B. M., Oksvold, P., Kampf, C., Djureinovic, D., Odeberg, J., Habuka, M., Tahmasebpoor, S., Danielsson, A., Edlund, K., Asplund, A., Sjöstedt, E., Lundberg, E., Szigyarto, C. A.-K., Skogs, M., Takanen, J. O., Berling, H., Tegel, H., Mulder, J., Nilsson, P., Schwenk, J. M., Lindskog, C., Danielsson, F., Mardinoglu, A., Sivertsson, A., Feilitzen, von, K., Forsberg, M., Zwahlen, M., Olsson, I., Navani, S., Huss, M., Nielsen, J., Ponten, F., Uhlén, M., 2014. Analysis of the human tissue-specific expression by genome-wide integration of transcriptomics and antibody-based proteomics. Mol. Cell Proteomics 13, 397–406. doi:10.1074/mcp.M113.035600

Finkbeiner, S. R., Hill, D. R., Altheim, C. H., Dedhia, P. H., Taylor, M. J., Tsai, Y.-H., Chin, A. M., Mahe, M. M., Watson, C. L., Freeman, J. J., Nattiv, R., Thomson, M., Klein, O. D., Shroyer, N. F., Helmrath, M. A., Teitelbaum, D. H., Dempsey, P. J., Spence, J. R., 2015. Transcriptome-wide Analysis Reveals Hallmarks of Human Intestine Development and Maturation In Vitro and In Vivo. Stem Cell Reports 4, 1140–1155. doi:10.1016/j.stemcr.2015.04.010

Firth, A. L., Dargitz, C. T., Qualls, S. J., Menon, T., Wright, R., Singer, O., Gage, F. H., Khanna, A., Verma, I. M., 2014. Generation of multiciliated cells in functional airway epithelia from human induced pluripotent stem cells. Proceedings of the National Academy of Sciences 111, E1723–30. doi: 10.1073/pnas.1403470111

Ghaedi, M., Calle, E. A., Mendez, J. J., Gard, A. L., Balestrini, J., Booth, A., Bove, P. F., Gui, L., White, E. S., Niklason, L. E., 2013. Human iPS cell-derived alveolar epithelium repopulates lung extracellular matrix. J. Clin. Invest. 123, 4950–4962. doi:10.1172/JCI68793

Ghosh, M., Ahmad, S., White, C. W., Reynolds, S. D., 2017. Transplantation of Airway Epithelial Stem/Progenitor Cells: A Future for Cell-Based Therapy. American Journal of Respiratory Cell and Molecular Biology 56, 1–10. doi: 10.1165/rcmb.2016-0181MA

Gilpin, S. E., Ren, X., Okamoto, T., Guyette, J. P., Mou, H., Rajagopal, J., Mathisen, D. J., Vacanti, J. P., Ott, H. C., 2014. Enhanced lung epithelial specification of human induced pluripotent stem cells on decellularized lung matrix. Ann. Thorac. Surg. 98, 1721–9–discussion 1729. doi: 10.1016/j.athoracsur.2014.05.080

Goss, A. M., Tian, Y., Tsukiyama, T., Cohen, E. D., Zhou, D., Lu, M. M., Yamaguchi, T. P., Morrisey, E. E., 2009. Wnt2/2b and & beta;-Catenin Signaling Are Necessary and Sufficient to Specify Lung Progenitors in the Foregut. Developmental Cell 17, 290–298. doi:10.1016/j.devcel.2009.06.005

Gotoh, S., Ito, I., Nagasaki, T., Yamamoto, Y., Konishi, S., Korogi, Y., Matsumoto, H., Muro, S., Hirai, T., Funato, M., Mae, S.-I., Toyoda, T., Sato-Otsubo, A., Ogawa, S., Osafune, K., Mishima, M., 2014. Generation of alveolar epithelial spheroids via isolated progenitor cells from human pluripotent stem cells. Stem Cell Reports 3, 394–403. doi:10.1016/j.stemcr.2014.07.005

Green, M. D., Chen, A., Nostro, M.-C., d’Souza, S. L., Schaniel, C., Lemischka, I. R., Gouon-Evans, V., Keller, G., Snoeck, H.-W., 2011. Generation of anterior foregut endoderm from human embryonic and induced pluripotent stem cells. Nat Biotechnol 1–7. doi:10.1038/nbt.1788

Harris-Johnson, K. S., Domyan, E. T., Vezina, C. M., Sun, X., 2009. beta-Catenin promotes respiratory progenitor identity in mouse foregut. Proceedings of the National Academy of Sciences 106, 16287–16292. doi: 10.1073/pnas.0902274106

Herriges, J. C., Verheyden, J. M., Zhang, Z., Sui, P., Zhang, Y., Anderson, M. J., Swing, D. A., Zhang, Y., Lewandoski, M., Sun, X., 2015. FGF-Regulated ETV Transcription Factors Control FGF-SHH Feedback Loop in Lung Branching. Developmental Cell 35, 322–332. doi:10.1016/j.devcel.2015.10.006

Hines, E. A., Sun, X., 2014. Tissue crosstalk in lung development. J Cell Biochem 115, 1469–1477. doi:10.1002/jcb.24811

Huang, S. X.L., Islam, M. N., O’Neill, J., Hu, Z., Yang, Y.-G., Chen, Y.-W., Mumau, M., Green, M. D., Vunjak-Novakovic, G., Bhattacharya, J., Snoeck, H.-W., 2013. efficient generation of lung and airway epithelial cells from human pluripotent stem cells. Nat Biotechnol 1–11. doi:10.1038/nbt.2754

Kim, H. Y., Nelson, C. M., 2012. Extracellular matrix and cytoskeletal dynamics during branching morphogenesis. Organogenesis 8, 56–64. doi:10.4161/org.19813

Konishi, S., Gotoh, S., Tateishi, K., Yamamoto, Y., Korogi, Y., Nagasaki, T., Matsumoto, H., Muro, S., Hirai, T., Ito, I., Tsukita, S., Mishima, M., 2015. Directed Induction of Functional Multi-ciliated Cells in Proximal Airway Epithelial Spheroids from Human Pluripotent Stem Cells. Stem Cell Reports 0. doi:10.1016/j.stemcr.2015.11.010

Kopp, J. L., Dubois, C. L., Schaffer, A. E., Hao, E., Shih, H. P., Seymour, P. A., Ma, J., Sander, M., 2011. Sox9+ ductal cells are multipotent progenitors throughout development but do not produce new endocrine cells in the normal or injured adult pancreas. Development 138, 653–665. doi:10.1242/dev.056499

Lange, A. W., Sridharan, A., Xu, Y., Stripp, B. R., Perl, A.-K., Whitsett, J. A., 2015. Hippo/Yap signaling controls epithelial progenitor cell proliferation and differentiation in the embryonic and adult lung. Journal of Molecular Cell Biology 7, 35–47. doi:10.1093/jmcb/mju046

Longmire, T. A., Ikonomou, L., Hawkins, F., Christodoulou, C., Cao, Y., Jean, J. C., Kwok, L. W., Mou, H., Rajagopal, J., Shen, S. S., Dowton, A. A., Serra, M., Weiss, D. J., Green, M. D., Snoeck, H.-W., Ramirez, M. I., Kotton, D. N., 2012. Efficient derivation of purified lung and thyroid progenitors from embryonic stem cells. Cell Stem Cell 10, 398–411. doi:10.1016/j.stem.2012.01.019

Love, M. I., Huber, W., Anders, S., 2014. Moderated estimation of fold change and dispersion for RNA-seq data with DESeq2. Genome Biol. 15, 31–21. doi: 10.1186/s13059-014-0550-8

Lu, B. C., Cebrian, C., Chi, X., Kuure, S., Kuo, R., Bates, C. M., Arber, S., Hassell, J., MacNeil, L., Hoshi, M., Jain, S., Asai, N., Takahashi, M., Schmidt-Ott, K. M., Barasch, J., D’Agati, V., Costantini, F., 2009. Etv4 and Etv5 are required downstream of GDNF and Ret for kidney branching morphogenesis. Nat Genet 41, 1295–1302. doi:10.1038/ng.476

Mahoney, J. E., Mori, M., Szymaniak, A. D., Varelas, X., Cardoso, W. V., 2014. The Hippo Pathway Effector Yap Controls Patterning and Differentiation of Airway Epithelial Progenitors. Dev. Cell 1–14. doi:10.1016/j.devcel.2014.06.003

McCauley, K. B., Hawkins, F., Serra, M., Thomas, D. C., Jacob, A., Kotton, D. N., 2017. Efficient Derivation of Functional Human Airway Epithelium from Pluripotent Stem Cells via Temporal Regulation of Wnt Signaling. Cell Stem Cell. doi:10.1016/j.stem.2017.03.001

Metzger, R. J., Klein, O. D., Martin, G. R., Krasnow, M. A., 2008. The branching programme of mouse lung development. Nature 453, 745–750. doi: 10.1038/Nature07005

Miller, A. J., Spence, J. R., 2017. In Vitro Models to Study Human Lung Development, Disease and Homeostasis. Physiology (Bethesda) 32, 246–260. doi: 10.1152/physiol.00041.2016

Moens, C. B., Auerbach, A. B., Conlon, R. A., Joyner, A. L., Rossant, J., 1992. A targeted mutation reveals a role for N-myc in branching morphogenesis in the embryonic mouse lung. Genes Dev. 6, 691–704. doi:10.1101/gad.6.5.691

Morrisey, E. E., Cardoso, W. V., Lane, R. H., Rabinovitch, M., Abman, S. H., Ai, X., Albertine, K. H., Bland, R. D., Chapman, H. A., Checkley, W., Epstein, J. A., Kintner, C. R., Kumar, M., Minoo, P., Mariani, T. J., McDonald, D. M., Mukouyama, Y.-S., Prince, L. S., Reese, J., Rossant, J., Shi, W., Sun, X., Werb, Z., Whitsett, J. A., Gail, D., Blaisdell, C. J., Lin, Q. S., 2013. Molecular determinants of lung development. Ann Am Thorac Soc 10, S12–6. doi: 10.1513/AnnalsATS.201207-036OT

Morrisey, E. E., Hogan, B. L.M., 2010. Preparing for the First Breath: Genetic and Cellular Mechanisms in Lung Development. Developmental Cell 18, 8–23. doi:10.1016/j.devcel.2009.12.010

Motoyama, J., Liu, J., Mo, R., Ding, Q., Post, M., Hui, C. C., 1998. Essential function of Gli2 and Gli3 in the formation of lung, trachea and oesophagus. Nat Genet 20, 54–57. doi:10.1038/1711

Nikolić, M. Z., Caritg, O., Jeng, Q., Johnson, J.-A., Sun, D., Howell, K. J., Brady, J. L., Laresgoiti, U., Allen, G., Butler, R., Zilbauer, M., Giangreco, A., Rawlins, E. L., 2017. Human embryonic lung epithelial tips are multipotent progenitors that can be expanded in vitro as long-term self-renewing organoids. Elife 6, e26575. doi:10.7554/eLife.26575

Okubo, T., Knoepfler, P. S., Eisenman, R. N., Hogan, B. L.M., 2005. Nmyc plays an essential role during lung development as a dosage-sensitive regulator of progenitor cell proliferation and differentiation. Development 132, 1363–1374. doi:10.1242/dev.01678

Ornitz, D. M., Yin, Y., 2012. Signaling Networks Regulating Development of the Lower Respiratory Tract. Cold Spring Harb Perspect Biol 4, a008318–a008318. doi:10.1101/cshperspect.a008318

Perl, A.-K.T., Kist, R., Shan, Z., Scherer, G., Whitsett, J. A., 2005. Normal lung development and function after Sox9 inactivation in the respiratory epithelium. genesis 41, 23–32. doi:10.1002/gene.20093

Rawlins, E. L., 2010. The building blocks of mammalian lung development. Dev. Dyn. 240, 463–476. doi:10.1002/dvdy.22482

Rawlins, E. L., Clark, C. P., Xue, Y., Hogan, B. L.M., 2009. The Id2+ distal tip lung epithelium contains individual multipotent embryonic progenitor cells. Development 136, 3741–3745. doi:10.1242/dev.037317

Rock, J. R., Hogan, B. L.M., 2011. Epithelial Progenitor Cells in Lung Development, Maintenance, Repair, and Disease. Annu. Rev. Cell Dev. Biol. 27, 493–512. doi:10.1146/annurev-cellbio-100109-104040

Rockich, B. E., Hrycaj, S. M., Shih, H.-P., Nagy, M. S., Ferguson, M. A.H., Kopp, J. L., Sander, M., Wellik, D. M., Spence, J. R., 2013. Sox9 plays multiple roles in the lung epithelium during branching morphogenesis. Proceedings of the National Academy of Sciences. doi: 10.1073/pnas.1311847110

Sekine, K., Ohuchi, H., Fujiwara, M., Yamasaki, M., Yoshizawa, T., Sato, T., Yagishita, N., Matsui, D., Koga, Y., Itoh, N., Kato, S., 1999. Fgf10 is essential for limb and lung formation. Nat Genet 21, 138–141. doi:10.1038/5096

Shu, W., Guttentag, S., Wang, Z., Andl, T., Ballard, P., Lu, M. M., Piccolo, S., Birchmeier, W., Whitsett, J. A., Millar, S. E., Morrisey, E. E., 2005. Wnt/beta-catenin signaling acts upstream of N-myc, BMP4, and FGF signaling to regulate proximal-distal patterning in the lung. Dev Biol 283, 226–239.

Spence, J. R., Lange, A. W., Lin, S.-C.J., Kaestner, K. H., Lowy, A. M., Kim, I., Whitsett, J. A., Wells, J. M., 2009. Sox17 Regulates Organ Lineage Segregation of Ventral Foregut Progenitor Cells. Developmental Cell 17, 62–74. doi:10.1016/j.devcel.2009.05.012

Spence, J. R., Mayhew, C. N., Rankin, S. A., Kuhar, M. F., Vallance, J. E., Tolle, K., Hoskins, E. E., Kalinichenko, V. V., Wells, S. I., Zorn, A. M., Shroyer, N. F., Wells, J. M., 2011. Directed differentiation of human pluripotent stem cells into intestinal tissue in vitro. Nature 470, 105–109. doi:10.1038/nature09691

Stahlman, M. T., Gray, M. P., Falconieri, M. W., Whitsett, J. A., Weaver, T. E., 2000. Lamellar body formation in normal and surfactant protein B-deficient fetal mice. Lab. Invest. 80, 395–403.

Tsai, Y.-H., Hill, D. R., Kumar, N., Huang, S., Chin, A. M., Dye, B. R., Nagy, M. S., Verzi, M. P., Spence, J. R., 2016. LGR4 and LGR5 Function Redundantly During Human Endoderm Differentiation. Cell Mol Gastroenterol Hepatol 2, 648–662.e8. doi: 10.1016/j.jcmgh.2016.06.002

Vanhecke, D., Herrmann, G., Graber, W., Hillmann-Marti, T., Mühlfeld, C., Studer, D., Ochs, M., 2010. Lamellar body ultrastructure revisited: high-pressure freezing and cryo-electron microscopy of vitreous sections. Histochem. Cell Biol. 134, 319–326. doi:10.1007/s00418-010-0736-4

Varner, V. D., Nelson, C. M., 2014. Cellular and physical mechanisms of branching morphogenesis. Development 141, 2750–2759. doi:10.1242/dev.104794

Weaver, M., Dunn, N. R., Hogan, B. L., 2000. Bmp4 and Fgf10 play opposing roles during lung bud morphogenesis. Development 127, 2695–2704.

White, A. C., Xu, J., Yin, Y., Smith, C., Schmid, G., Ornitz, D. M., 2006. FGF9 and SHH signaling coordinate lung growth and development through regulation of distinct mesenchymal domains. Development 133, 1507–1517. doi:10.1242/dev.02313

Wong, A. P., Bear, C. E., Chin, S., Pasceri, P., Thompson, T. O., Huan, L.-J., Ratjen, F., Ellis, J., Rossant, J., 2012. Directed differentiation of human pluripotent stem cells into mature airway epithelia expressing functional CFTR protein. Nat Biotechnol. doi:10.1038/nbt.2328

Yin, Y., Wang, F., Ornitz, D. M., 2011. Mesothelial-and epithelial-derived FGF9 have distinct functions in the regulation of lung development. Development 138, 3169–3177. doi:10.1242/dev.065110

Yin, Y., White, A. C., Huh, S.-H., Hilton, M. J., Kanazawa, H., Long, F., Ornitz, D. M., 2008. An FGF-WNT gene regulatory network controls lung mesenchyme development. Developmental Biology 319, 426–436. doi:10.1016/j.ydbio.2008.04.009

Zhang, Y., Yokoyama, S., Herriges, J. C., Zhang, Z., Young, R. E., Verheyden, J. M., Sun, X., 2016. E3 ubiquitin ligase RFWD2 controls lung branching through protein-level regulation of ETV transcription factors. Proceedings of the National Academy of Sciences 201603310. doi: 10.1073/pnas.1603310113

Zhao, R., Fallon, T. R., Saladi, S. V., Pardo-Saganta, A., Villoria, J., Mou, H., Vinarsky, V., Gonzalez-Celeiro, M., Nunna, N., Hariri, L. P., Camargo, F., Ellisen, L. W., Rajagopal, J., 2014. Yap tunes airway epithelial size and architecture by regulating the identity, maintenance, and self-renewal of stem cells. Developmental Cell 30, 151–165. doi:10.1016/j.devcel.2014.06.004

